# The complex evolutionary history of *Capparis* L. (Capparaceae Juss.) in Australia, and the description of five new species and two new subspecies

**DOI:** 10.1101/2025.03.06.641633

**Authors:** W.E. Cooper, D.M. Crayn, F.A. Zich, E.M. Joyce

## Abstract

*Capparis* comprises ∼145 species, of which ∼21 occur in Australia; however, the relationships and taxonomic status of Australian *Capparis* taxa remain to be tested. We present phylogenies of all Australian *Capparis* taxa by analysing Angiosperms353 loci using coalescent and concatenated approaches. All trees resolve *Capparis* as monophyletic, with a whole-genome duplication (WGD) event detected at the crown. *Capparis* sect. *Capparis* and sect. *Busbeckea* are monophyletic, but sect. *Monostichocalyx* is non-monophyletic. The relationships of species within sect. *Busbeckea* are poorly supported due to rapid radiation following ancient WGD. The relationships of taxa within sect. *Capparis* and the clades of sect. *Monostichocalyx* are well-supported, with some incomplete lineage sorting. *Capparis spinosa* is geographically, morphologically and phylogenetically structured across northern Australia. Based on these results, we describe five new species and two new subspecies of *Capparis*, bringing the total number of species in Australia to 26*. Capparis xylocarpa*, *C. megacarpa*, *C. loxophleba* and *C. splendidissima* are newly described. *Capparis loranthifolia* var. *bancroftii* is raised to species level as *C. bancroftii*. *Capparis spinosa* subsp. *nummularia* is split into three subspecies: subsp. *formicosa*, subsp. *insularis*, and subsp. *nummularia*. We provide a key for all Australian taxa.

## Introduction

*Capparis* L. (Capparaceae Juss.) is a genus of tropical and subtropical vines, shrubs and trees occurring in Europe, Africa, Asia, Australia, and Indian and Pacific Ocean islands (Cornejo 2018; Maurya *et al*. 2023). The best-known member of the genus is the Caper Bush *Capparis spinosa* L. — a species distributed from Europe throughout Asia, Australia and the Pacific Islands. Its buds (‘capers’) and fruits (‘caper berries’) have been consumed for millennia for their nutritional and medicinal value, and are an important economic crop for many Mediterranean countries (Hansen 1991; Romeo *et al*. 2007; Jiang *et al*. 2007). Despite the importance of the genus, the number of species in *Capparis* is uncertain, with 142–146 accepted species (Maurya *et al*. 2023; WFO 2024) and an additional 43 unplaced names (WFO 2024). Little molecular work has been done on the genus, with previous studies of the genus based on few chloroplast markers (Hall *et al*. 2002; Tamboli *et al*. 2018). The most recent and comprehensive phylogeny based on sequences of *matK*, *trnL-F* and *rbcL* sampled 43 species, including three species that occur in Australia (Maurya *et al*. 2023). As such, much phylogenetic and taxonomic work is needed to classify the diversity in this genus and understand its evolutionary history.

*Capparis* L. is one of three Capparaceae genera naturally occurring in Australia — the others being *Cadaba* Forssk. and *Crateva* L.. The Flora of Australia documents 18 species of *Capparis* in three sections (sect. *Monostichocalyx* Radlk., sect. *Capparis* and sect. *Busbeckea* (Endl.) Hook.f.), including 14 endemics (Hewson 1982, Green 1994; Fig. 1). Hewson’s (1982) revision of mainland species also included five ‘doubtful taxa’, noting that the limits of several taxa were in need of further study. Since then, an additional three endemic species have been recognised for Australia; *C. velutina* P.I.Forst. (Forster 1997), *C. batianoffii* Guymer (Guymer 2008), and *C. anomala* (F.Muell.) Byng & Christenh. (Christenhusz *et al*. 2018). *Capparis anomala* was transferred to *Capparis* from *Apophyllum* F.Muell. based on limited molecular phylogenetic evidence and no morphological study, and its generic placement requires verification (Christenhusz *et al*. 2018). This brought the total number of species accepted in Australia to 21. Of these, *C. loranthifolia* Lindl., *C. arborea* (F.Muell.) Maiden*, C. lucida* (DC.) R.Br. ex Benth. and *C. spinosa* have been in particular need of study to clarify taxon boundaries.

**Fig. 1.**
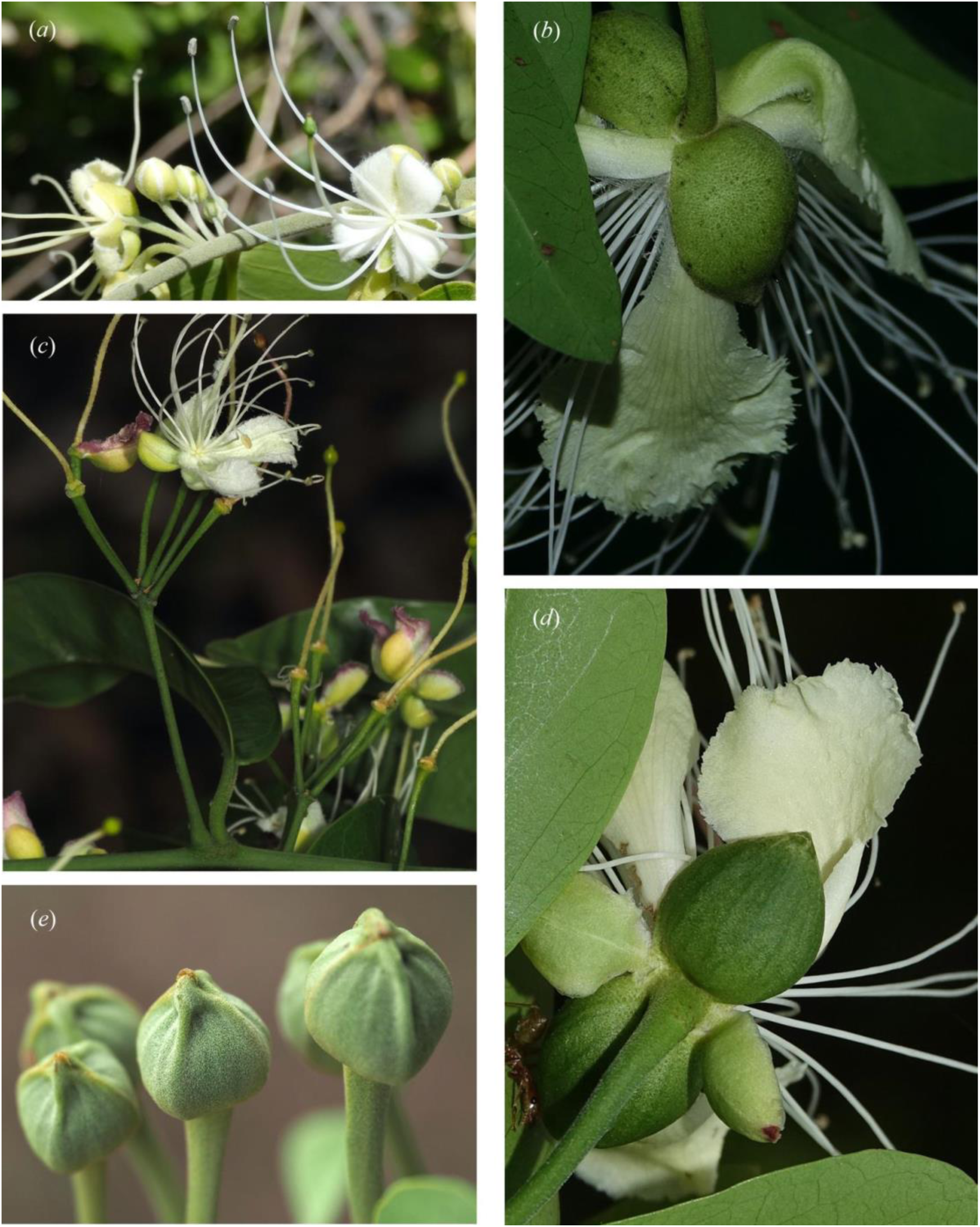
Various Australian *Capparis* inflorescences, buds and flowers: (*a*) *Capparis quiniflora* with supra-axillary solitary flowers or serial rows of up to 5, *Cooper 2600*. (*b*) *Capparis arborea* showing outer pair of sepals without ribs, inner pair of sepals have dehisced and fallen, *Cooper 2744*. (*c*) *Capparis lanceolaris* with pedicellate umbels, *Cooper 3116*. (*d*) *Capparis lucida* showing outer pair of sepals ribbed and inner pair of sepals smooth and with a different and thinner texture as is the case with all Australian *Capparis, Cooper 2616*. (*e*) *Capparis canescens* showing 4-ribbed buds, *Cooper 3123*. Photographs: (*a*), (*c*) & (*e*) W. Cooper, (*b*) & (*d*) R. Jensen.

*Capparis loranthifolia* is a widespread species endemic to Australia, currently subdivided into two varieties; var. *loranthifolia* and var. *bancroftii* C.T.White ex M.Jacobs. The fruits of the varieties have been considered distinct, and Hewson (1982) remarked that ‘Further study of this species may show that var. *bancroftii* should be raised to specific rank’. Additional morphological and molecular work is needed to verify the taxonomic status of these varieties.

*Capparis arborea* is distributed from central New South Wales to the Cape York Peninsula. It is morphologically variable and easily confused with *C. lucida* in some parts of its range, leading Hewson (1982) to suggest that further study was required to test the boundaries between these two species. Molecular data can be used to test the affinities of these species in individuals with overlapping morphological characters.

*Capparis spinosa* in Australia has a complex taxonomic history, variously attributed to *C. nummularia* DC., *C. spinosa* var. *nummularia* F.M.Bailey, *C. nummularia* DC. var. *nummularia*, *C. nummularia* var. *typica* Domin, *C. nummularia* var. *minor* Domin, and *C. spinosa* var. *mariana* (Jacq.) K.Schum. In the latest taxonomic work on the species, all these names were synonymised under *C. spinosa* subsp. *nummularia* (DC.) Fici (Fici 2003, 2015); however, the taxon is still treated as *C. nummularia* in Queensland (Queensland Government 2023; CHAH 2024). The convoluted taxonomic history reflects the morphological variation in disjunct parts of its range in Western Australia, the Northern Territory, central Queensland and the offshore islands of Queensland. Based on the observation of seedling leaf traits of individuals from the Northern Territory and Western Australia in an informal common garden study, Fici (2003) concluded that morphological differences were the result of environment, and that all morphotypes in Australia belonged to one subspecies. However, the genetic distinctiveness of these morphotypes has never been studied. Further complicating classification is the resemblance of the island form of *C. spinosa* subsp. *nummularia* to *C. spinosa* subsp. *cordifolia* (Lam.) Fici from New Guinea and the Pacific Islands. Therefore, whether these geographically disjunct morphotypes are genetically distinct, and whether they are distinct from *C. spinosa* var. *cordifolia*, required testing.

In addition to the 21 currently accepted species in Australia, two phrase-named taxa — *C*. sp. Bamaga (V.Scarth-Johnson 1048A) Qld Herbarium and *C*. sp. Coen (L.S.Smith 11862) Qld Herbarium — are recognised by the Australian Plant Census as putatively new (CHAH 2024). Recent fieldwork by the first author has identified two additional putative new species: one with affinities to *C. ornans* F.Muell. ex Benth. but with distinct morphology and a disjunct, rainforest distribution (herein referred to as ‘*Capparis* sp. Wet Tropics vine’); and one novel entity from the Maleny region of Queensland (herein referred to as ‘*Capparis* sp. hairy-leaved form’). Phylogenetic work and morphological study is needed to ascertain the affinities of these putative new species and phrase-named species, and whether they warrant recognition as species.

In this study, we generated a next-generation sequencing phylogeny of all Australian *Capparis* taxa to reconstruct their evolutionary history and test species boundaries. Specifically, we aimed to (1) infer the relationships and evolutionary history of all Australian *Capparis* species; (2) test the monophyly of morphologically variable species; (3) test the genetic distinctiveness of *C. spinosa* subsp. *nummularia* across its range and its relationship to *C. spinosa* subsp. *cordifolia;* (4) assess whether any entities within *Capparis* warrant a change in taxonomic rank or circumscription, and (5) provide a diagnosis for new and recircumscribed taxa, and a key for all Australian species.

## Materials and methods

### Sampling

A total of 44 *Capparis* samples were obtained for this study, including all Australian *Capparis* taxa, the two phrase-named taxa, the putative novelties, 15 samples of *C. spinosa* subsp. *nummularia* across its geographic range and one sample of a non-Australian species – *C. spinosa* subsp. *cordifolia* (Supplementary material 1). Of the 15 *C. spinosa* subsp. *nummularia* samples included, four were from Western Australia (WA), five were from inland Queensland (QLD), four were from islands off the east coast of Queensland (E. QLD), and two were from the Northern Territory (NT). This sampling allowed us to assess the monophyly of *Capparis spinosa* subsp. *nummularia*, and the relationship of the insular ‘E. QLD’ morphotype of *C. spinosa* subsp. *nummularia* to *C. spinosa* subsp. *cordifolia* from Papua New Guinea and the Pacific Islands and mainland Australian morphotypes of *C. spinosa* subsp. *nummularia*. It also allowed us to assess whether the geographically disjunct populations of *C. spinosa* subsp. *nummularia* are genetically distinct and warrant recognition as separate taxa. Three samples of *C. arborea* — two from the northern morphotype and one from the southern morphotype — were included to assess the monophyly of the species. Both subspecies of *C. loranthifolia* were sampled to determine whether they are monophyletic. *Buchholzia coriacea* Engl., *Cadaba capparoides* DC., *Maerua glauca* Chiov., *Thilachium africanum* Lour. and *Boscia albitrunca* Gilg & Gilg-Ben. were included as outgroups based on published phylogenies (Hall *et al*. 2002; Maurya *et al*. 2023; Zuntini *et al*. 2024).

All *Capparis* samples were derived from leaf material that was either collected in the field and dried on silica gel, or sampled from herbaria. Herbarium material was taken from sheets at BRI, CANB, CNS, M and PERTH (Supplementary Material 1). Data for outgroup samples were sourced from the Sequence Read Archive (SRA) via the Kew Tree of Life Explorer (Royal Botanic Gardens, Kew 2024).

### DNA extraction

For all *Capparis* samples, genomic DNA was extracted from 40 mg of dried leaf material using a modified CTAB protocol (Doyle and Doyle 1987). At the isopropanol precipitation step, samples were left to precipitate at −20℃ for 24 hours. DNA was eluted in 60 µl, and quality and quantity was ascertained using a NanoDrop 1000 Spectrophotometer (Thermo Fisher Scientific, Massachusetts, USA) and Quantus Fluorometer (Promega Corporation, Wisconsin, USA).

### Library preparation

Following DNA extraction, all *Capparis* samples were prepared for target capture sequencing using the Angiosperm353 bait kit, with the exception of five samples: *C. spinosa* subsp. *nummularia* 9, *C. spinosa* subsp. *nummularia* 10, *C*. sp. hairy-leaved form, *C. umbonata* Lindl. and *C. thozetiana F.Muell*. These samples either failed the initial round of sequencing or were added at a later date, and genome skimming was performed on these samples because it was more economical.

When necessary, DNA samples were fragmented enzymatically. The NEBNext Ultra II FS Library Prep Kit (New England Biolabs, Ipswich, MA, USA) was used for all samples following the manufacturer’s instructions to generate libraries of approximately 350 bp. Libraries were combined into pools of 12–16, and pooled libraries were enriched using the Angiosperms353 probe kit (Johnson *et al*. 2019) by hybridising at 65°C with the Arbor Biosciences MyBaits Expert Plant Angiosperms353 v1 bait set with V5 chemistry (Cat. # 308108.v5). The hybridisation step was skipped for the five genome skimming libraries produced at a later date. Sequencing of Angiosperm353-enriched libraries was performed on a NovaSeq 6000 (Illumina Inc., San Diego, USA) at the Australian Genome Research Facility (Melbourne) with v1.5 chemistry and 150 bp paired-end reads. Sequencing of genome skimming samples was performed on a NovaSeq X (Illumina Inc., San Diego, USA) with 150 bp paired-end reads by Novogene (Munich, Germany).

### Species tree inference

All data was filtered for adapter sequences, poor quality base calls (with PHRED scores <13) and poor quality reads (with a minimum average PHRED score of 16) using the ‘clean’ command of CAPTUS (Ortiz *et al*. 2023). Angiosperm353 loci were then assembled from the clean data using the CAPTUS pipeline (Ortiz *et al*. 2023). *De novo* assembly was performed with the ‘assemble’ command under ‘CAPSKIM’ settings. Locus extraction was conducted with the ‘extract’ command against the mega target file of McLay *et al*. (2021) filtered to Brassicales sequences. Prior to locus extraction, the filtered target file was checked and fixed using the ‘check_targetfile’ and ‘fix_targetfile’ commands of HybPiper (Johnson *et al*. 2016). Only sequences that had a minimum percentage identity (‘--nuc_min_score’) of 75% were retained, and all paralogs were retained. The ‘align’ command of CAPTUS was then used to collect all sequences for each locus, including all paralogs, without aligning or cleaning them (‘—collect_only’).

To effectively deal with paralogs and infer orthologs (Morales-Briones *et al*. 2021; Joyce *et al*. 2024), exon sequences of each locus were processed through Paragone (Jackson *et al*. 2023). In this pipeline, all sequences were aligned and trimmed (command ‘check_and_align’), and the trimmed alignments were used to generate raw homolog trees (i.e., gene trees including all paralogs; command ‘alignment_to_tree’). Raw homolog trees were then visually inspected and cleaned for monophyletic paralog tips, long branches (--treeshrink_q_value = 0.10), and deep paralogs (--cut_deep_paralogs_internal_branch_length_cutoff = 0.8). Using the command ‘align_selected_and_tree’, the sequences remaining in the tree were then realigned and trimmed (--trimal_gapthreshold = 0.25), and clean homolog trees were generated. From the clean homolog trees, orthologs were identified with the ‘prune_paralogs’ command using the Monophyletic Outgroups algorithm of Yang & Smith (2014). Inferred orthologs were then extracted, aligned and trimmed using the command ‘final_alignments’.

Due to the high number of paralogs in the dataset, two approaches to inferring trees were used. The first used the paralogs in our homolog trees to inform species tree inference with a quartet-based method (our ‘paralog-informed’ species tree), and the second used inferred orthologs to estimate species trees using coalescent and concatenated Maximum Likelihood methods (our ‘coalescent ortholog’ and ‘concatenated ortholog’ species trees, respectively). For the first approach, the cleaned homolog trees for all loci were used to estimate a paralog-informed species tree using ASTRAL-Pro v1.16.2.4 (Zhang *et al*. 2020). For our ortholog species trees, ortholog alignments of each locus were concatenated with AMAS (Borowiec 2016), and the ortholog concatenated species tree was estimated in IQ-TREE 2 from the concatenated alignment (Nguyen *et al*. 2015; Minh *et al*. 2020). The appropriate substitution model was chosen using ModelFinder Plus option (–MFP), and support was estimated using 1000 ultrafast bootstrap replicates (Lanfear *et al*. 2012; Kalyaanamoorthy *et al*. 2017; Hoang *et al*. 2018). For the ortholog coalescent species tree, ortholog gene trees were estimated from the ortholog alignments using IQ-TREE 2 with the appropriate substitution model chosen using ModelFinder, and 1000 ultrafast bootstrap replicates (Lanfear *et al*. 2012; Nguyen *et al*. 2015; Kalyaanamoorthy *et al*. 2017; Hoang *et al*. 2018; Minh *et al*. 2020). Poorly supported branches in the resulting gene trees (BS <50) were collapsed using Newick Utils v1.6, (Junier & Zdobnov 2010). The cleaned ortholog gene trees were then used to infer a species tree using ASTRAL v5.7.8 (Mirarab *et al*. 2014).

### Investigation of phylogenetic conflict

To investigate phylogenetic conflict in the species trees, phyparts (Smith 2024) was used to compare ortholog gene tree topologies to the species tree. The scripts of Morales-Briones *et al*. (2021) were used to modify the output of phyparts to account for missing taxa and retrieve values of dis/concordance that are proportional to the number of informative nodes.

To investigate whether polyploidisation might play a role in explaining patterns of paralogy and phylogenetic conflict in Australian *Capparis*, we used HybPhaser (Nauheimer et al. 2021) to characterise average heterozygosity and allele divergence. Heterozygosity and allele divergence has been demonstrated to be an effective indicator of ploidy level in the closely-related family Brassicaceae (Hendriks *et al*. 2023; Walden *et al*. 2024). As such, we used HybPhaser to test for signal of ploidy variation in the samples of our dataset. As HybPhaser is dependent on the output of HybPiper, we assembled exon sequences from the dataset cleaned with CAPTUS using the HybPiper v2 ‘assemble’ command (Johnson *et al*. 2016). Trimmed reads were mapped against the filtered nucleotide mega target file using BWA. The ‘1_generate_consensus_sequences’ script of HybPhaser v2 (Nauheimer *et al*. 2021) was then used to remap the reads of the HybPiper output. The HybPhaser R script ‘1a_count_snps.R’ was run to identify and summarise the SNPs from alleles and paralogs for each sample, and ‘1b_assess_dataset.R’ was used to calculate heterozygosity at different thresholds, and allele divergence. Summary statistics of gene recovery and proportion of SNPs were generated for all samples and loci.

To further assess our dataset for polyploidisation events and to locate their potential occurrence in the phylogeny, we used our homolog trees to map duplication events on our species tree. To do this, we followed the whole-genome duplication (WGD) mapping algorithm of Yang *et al*. (2018). In brief, we trimmed all our clean homolog trees to ‘orthogroups’, i.e. subtrees of rooted ingroup samples representing the relationships of single copies within each homolog tree. Each orthogroup with an average bootstrap value above 50 was then compared to the species tree. If two or more taxa overlapped, a gene duplication event was counted at the most recent common ancestor of the species tree, with a maximum of one gene duplication event recorded per orthogroup. The proportion of orthogroups indicating a gene duplication event can then be calculated for each branch of the species tree, and outlying high values are indicative of branches where a WGD event is likely to have occurred.

### Morphological study

Morphology of Australian *Capparis* was assessed through examination of herbarium material from BRI, CNS, MEL, NSW and PERTH, as well as specimen images from K and L available online (Global Plants on JSTOR; https://plants.jstor.org/), and field observations. Particular attention was given to undescribed species, putative novelties and the morphologically variable species (*C*. *spinosa* subsp. *nummularia*, *C. arborea* and *C*. *loranthifolia*). Measurements of floral parts and fruits are based on material preserved in 70% ethanol as well as fresh material from the field.

## Results

### Locus extraction

An average of 342 (199–352) loci per sample were retrieved using CAPTUS, with a mean target recovery (across all copies of all genes) of 81.4% (51.5–91.2%). A high number of paralogs were extracted, with a mean of 1.6 (1.1–5.2) copies of each locus retrieved for each sample, and up to 42 copies retrieved of individual loci. ‘*Capparis* sp. Wet Tropics vine’ *C. shanesiana* F.Muell., *C. arborea* 2, *C. jacobsii* Hewson and *C. loranthifolia* var. *loranthifolia* had a particularly high number of paralogs, with an average of 2.9-5.2 copies per locus (Supplementary Material 2).

### Species tree estimation

Processing of our dataset through the Paragone pipeline retained 320 clean homolog trees. These were used to estimate our paralog-informed species tree using ASTRAL-Pro (Fig. 2). A total of 322 orthologs were inferred from our dataset, and all 322 ortholog gene trees were used to estimate the ortholog coalescent species tree with ASTRAL (Supplementary Material 3). For our ortholog concatenated species tree, input concatenated alignment comprised 322 loci of 235909 bp and 34101 parsimony-informative sites. The optimal substitution model for each locus was chosen with the ModelFinder function of IQ-TREE 2 (LnL = −934734, df=1607), and after 153 tree search iterations IQ-TREE 2 estimated a consensus tree with a log-likelihood of −934622 (Supplementary Material 4).

**Fig. 2.**
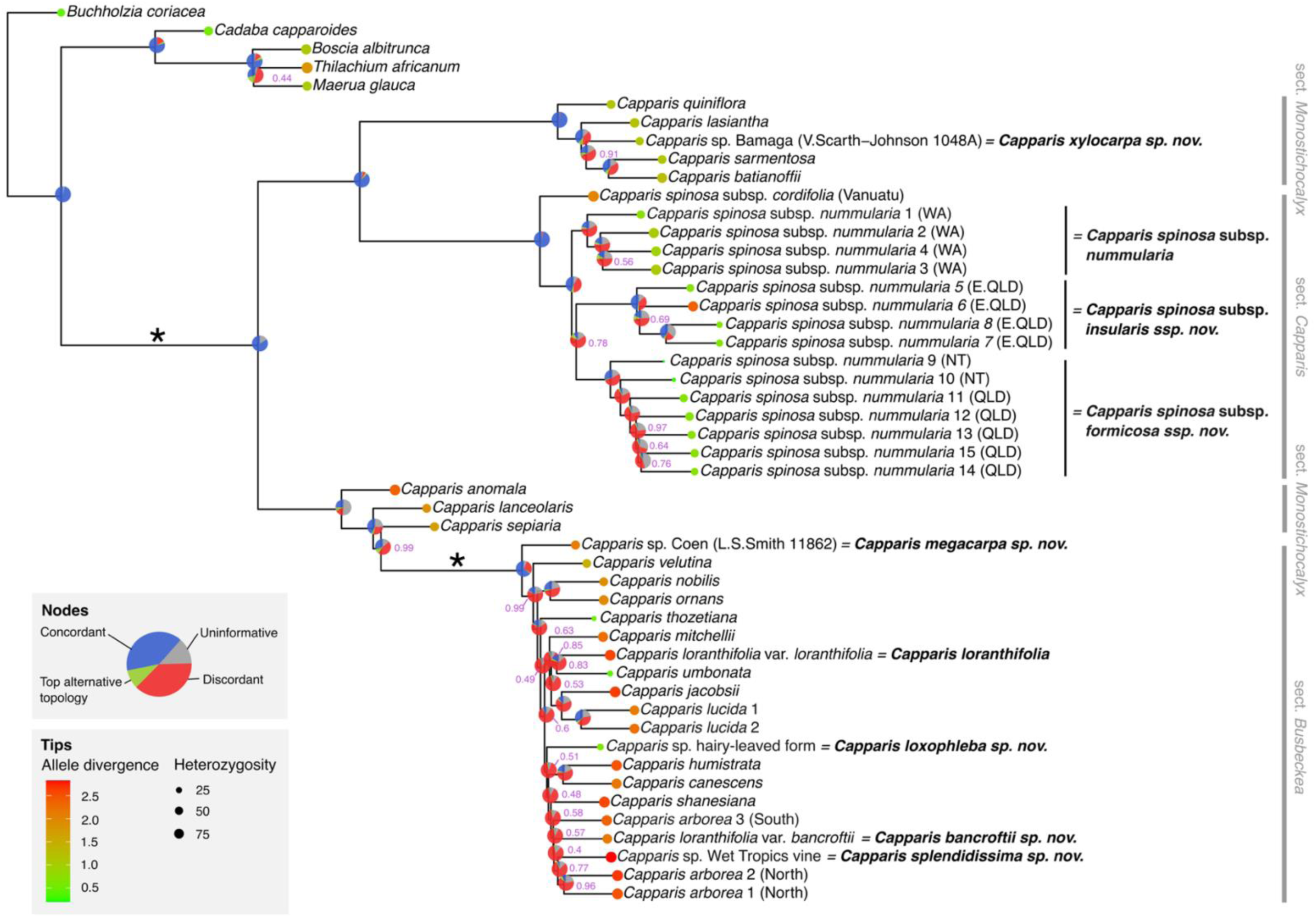
Paralog-informed coalescent tree of all Australian *Capparis* taxa, and *C. spinosa* subsp. *cordifolia* from Vanuatu, based on nuclear homolog trees analysed with ASTRAL-Pro. All forms of *C. spinosa* subsp. *nummularia* are denoted in brackets (see Figure 1). Currently recognised sections are indicated in grey to the right of the tree. Circles on tips indicate level of paralogy as estimated with HybPhaser; colour denotes mean allele divergence, and circle size indicates mean heterozygosity across all loci. Posterior probabilities for all nodes with a value <1.0 is indicated in purple; nodes without purple values received maximum support. Pie charts summarise the concordance of gene tree topology compared to species tree topology as calculated with Phyparts, with blue indicating the proportion of loci with a concordant topology for that node, green indicating the proportion of loci that support the top alternative topology, red denoting the proportion of loci that support conflicting topologies, and grey representing the proportion of uninformative loci for that node. Asterisks indicate the two branches where whole genome duplication events are inferred to have occurred based on WGD mapping of orthogroups.

Each species tree recovered three major clades in the same topology (Fig. 2; Supplementary Material 3-4). All samples of *Capparis spinosa*, including the Australian *C. spinosa* subsp. *nummularia* and the sample of *C. spinosa* subsp. *cordifolia* were monophyletic, corresponding to *C.* sect. *Capparis* (Fig. 2; Supplementary Material 3–4). This clade is always sister to a clade containing *C. quiniflora* DC., *C. lasiantha* R.Br. ex DC., *C*. sp. Bamaga (V.Scarth-Johnson 1048A), *C. sarmentosa* A.Cunn. ex Benth., and *C. batianoffii* – all of which (with the exception of *C*. sp. Bamaga (V.Scarth-Johnson 1048A)) belong to *C.* sect. *Monostichocalyx* (Fig. 2; Supplementary Material 3–4). In all species trees, this clade pair is sister to another major clade. This third major clade is comprised of a grade of species from *C.* sect. *Monostichocalyx* (*C. anomala, C. lanceolaris* DC. and *C. sepiaria* L.), which terminates in a clade containing 20 samples of taxa from *C.* sect. *Busbeckea* (Fig. 2; Supplementary Material 3–4). In all trees, the representatives of *C.* sect. *Busbeckea* form a clade with short branches, while the grade of *C.* sect. *Monostichocalyx* samples are more distantly related.

While the topology of deeper nodes was consistent across species trees, the topology towards the tips differed for some clades. Given that the paralog-informed analysis produced a topology that is supported by morphology, we show these results in Fig. 2. However, in cases where topology differs, these results are highlighted, and the relevant species trees can be seen in Supplementary Material 3–4.

In all species trees, the topology of the first major clade containing *Capparis quiniflora*, *C. lasiantha*, *C*. sp. Bamaga (V.Scarth-Johnson 1048A), *C. sarmentosa*, and *C. batianoffii* was consistent and well supported, with relatively little phylogenetic discordance (BS=99–100, PP=0.91–1.0) (Fig. 2, Supplementary Material 3–4).

In the second major clade containing *Capparis spinosa*, *C. spinosa* subsp. *cordifolia* is consistently retrieved as sister to a clade of *C. spinosa* subsp. *nummularia* (Fig. 2, Supplementary Material 3–4). Within the clade of *C. spinosa* subsp. *nummularia*, the Western Australian (WA) individuals are always monophyletic, as are the island form from eastern Queensland (E. QLD), and the Northern Territory (NT) individuals form a clade with individuals from inland Queensland (QLD) (Fig. 2). However, the topology of these three clades differs across the species trees; in the paralog-informed and concatenated ortholog species trees, the WA clade is sister to an (E. QLD:NT-QLD) clade (Fig. 2, Supplementary Material 4), while in the coalescent ortholog tree, the E. QLD clade is sister to a (WA: NT-QLD) clade (Supplementary Material 3). Across species trees, this topological discrepancy seems to be related to the relatively uncertain relationship of the NT-QLD clade with other *C. spinosa* subsp. *nummularia* clades. The stem node of the NT-QLD clade received moderate support (PP=0.75–0.78) in the paralog-informed and coalescent ortholog species trees (Fig. 2, Supplementary Material 3), high support (BS=0.99) in the concatenated ortholog species tree (Supplementary Material 4), and shows a high degree of phylogenetic discordance in the phyparts analysis (Fig. 2).

In the third major clade, the samples of *Capparis* sect. *Monostichocalyx* are always placed with the same topology with high support, with *C. anomala* being the earliest branching species in the grade, followed by *C. lanceolaris* and then *C. sepiaria*. However, the relationships of the samples in the *C.* sect. *Busbeckea* subclade differ across trees, mostly receiving poor support and having a very high degree of phylogenetic conflict throughout. What is clear is that in all analyses, *C.* sp. Coen (L.S.Smith 11862) is the earliest diverging lineage in the *C.* sect. *Busbeckea* subclade, with maximum support and little signal of phylogenetic conflict. *Capparis lucida* is consistently retrieved as monophyletic with maximum support and relatively low phylogenetic conflict. *Capparis arborea* is estimated as non-monophyletic in all species trees, with the northern individuals forming a clade and the southern sample of *C. arborea* always placed separately with poor support and high discordance. *Capparis loranthifolia* is also non-monophyletic in all species trees, with *C. loranthifolia* var. *loranthifolia* placed in a separate clade from *C. loranthifolia* var. *bancroftii* with low to moderate support and a high degree of discordance. The placement of the putative novelties ‘*C*. sp. Wet Tropics vine’ and ‘*C*. sp. hairy-leaved form’ differs across species trees. ‘*Capparis* sp. Wet Tropics vine’ is retrieved with the northern individuals of *C. arborea* in the paralog-informed and concatenated ortholog species tree with moderate support (PP=0.77, BS=70), but is placed as sister to a clade containing *C. humistrata* F.Muell., *C. canescens* G.Don, *C. arborea, C. shanesiana* and *C. loranthifolia* var. *bancroftii* in the coalescent ortholog analysis, with no support (PP=0.04). ‘*C*. sp. hairy-leaved form’ is always allied with *C. humistrata*, *C. canescens*, *C. shanesiana*, *C. arborea*, ‘*C*. sp. Wet Tropics vine’, and *C. loranthifolia* var. *bancroftii*, but its position within this clade differs across species trees.

### Ploidy characterisation and WGD mapping

After re-extracting the exon regions from the cleaned data with HybPiper, we retrieved an average of 292 loci per sample, with a mean recovery of 75%. We used the reads extracted by HybPiper for the HybPhaser pipeline to characterise heterozygosity and allele divergence (AD) of regions extracted for each sample, and gain an indication of variation in ploidy level. The mean AD for all extracted loci in Australian *Capparis* sampled was 1.36% (0.05–2.7%), and the average locus heterozygosity across all *Capparis* samples was 63.04% (6.15–94.96%) (Supplementary Material 2). Of the three major clades in Australian *Capparis*, the *C.* sect. *Capparis* clade had the lowest AD (0.74% [0.05–2.24]) and heterozygosity (51.45% [6.15– 92.47]), but with *C. spinosa* subsp. *nummularia* 6 and *C. spinosa* subsp. *cordifolia* (from Vanuatu) having higher than average values for the clade (Fig. 2; Supplementary Material 2). Its sister clade containing some samples of *C.* sect. *Monostichocalyx* had the next highest AD (1.03% [0.99–1.10]) and heterozygosity (62.28 [49.85–77.55]) (Fig. 2; Supplementary Material 2). The (*C.* sect. *Monostichocalyx* + *C.* sect. *Busbeckea*) clade showed the highest AD (1.87% [0.275–2.67]) and heterozygosity (71.28% [15.38–94.96]) overall, but with *C. thozetiana*, *C. umbonata* and *C*. *arborea* (hairy-leaved form) having lower values than average for the clade.

Our WGD mapping analysis showed some degree of gene duplication in 18 nodes across the tree, with most of these nodes having inferred duplication events in 0.5–9.9% of genes (Supplementary Material 5). An elevated gene duplication level (>1.5x the inter-quartile range) was mapped at two nodes: at the crown node of *Capparis* (32.1%) and at the crown node of *C.* sect. *Busbeckea* (29.2%) (Fig. 2, Supplementary Material 5), indicating that a WGD event has occurred at these two points of the phylogeny (Yang *et al*. 2018).

## Discussion

### The complex evolutionary history of *Capparis* in Australia

Our analysis shows that Australian *Capparis* is monophyletic, confirming the placement of *C. anomala* (transferred from *Apophyllum anomalum F.Muell.* in Christenhusz *et al*. 2018) within *Capparis*. This finding supports the results of previous molecular studies based on chloroplast and mitochondrial markers, where *Capparis s.s.* (Old World *Capparis* + *Apophyllum anomalum*) was found to be monophyletic (Hall *et al*. 2002; Cardinal-McTeague *et al*. 2016; Tamboli *et al*. 2018; Maurya *et al*. 2023). However, our findings are in conflict with the recent, angiosperm-wide phylogeny of Zuntini *et al*. (2024), where the three *Capparis* samples included were not monophyletic. This could have been caused by unbalanced sampling in the large phylogeny of Zuntini *et al*. (2024), or noise caused by paralogs in the nuclear dataset from WGD events common in Brassicales families (Cardinal-McTeague *et al*. 2016). On the other hand, it could also be an indication that when the full geographic range of *Capparis* is considered, rather than just Australian species as in our study, the genus is not monophyletic. Therefore, the monophyly of *Capparis* needs to be investigated further with a phylogenomic or transcriptomic dataset, enhanced sampling for the genus, and extensive outgroup sampling from other closely related genera in Capparaceae.

Our phylogeny indicates that there are three major clades of *Capparis* in Australia, but these clades do not correspond to current sectional classification of Australian species (Jacobs 1964; Hewson 1982). While *C*. sect *Busbeckea* and *C*. sect. *Capparis* are monophyletic, *C*. sect. *Monostichocalyx* is not monophyletic, as *C. lanceolaris* and *C. sepiaria* (of *C*. sect. *Monostichocalyx*) form a clade with *C. anomala* (section unassigned) and the *C*. sect. *Busbeckea* clade. This result is congruent with the findings of Maurya *et al*. (2023), who also retrieved a polyphyletic *C*. sect. *Monostichocalyx*. In line with our results, the authors found that *C. sepiaria* from the Philippines and *C. anomala* form a clade with representatives of other *C*. sect. *Monostichocalyx* species, and that *C*. sect *Busbeckea* was nested in this clade. Together, this clade is referred to by Maurya *et al*. 2023 as ‘*Capparis* Section I’, which was said to be characterised by its inflorescence structure of umbels, sub-umbels, corymbs or panicles. However, Maurya *et al*. (2023) did not include any of the other *C*. sect. *Monostichocalyx* species from Australia that form a separate clade (*C. quiniflora*, *C. lasiantha*, *C*. sp. Bamaga (V.Scarth-Johnson 1048A), *C. sarmentosa*, and *C. batianoffii*) in their phylogeny. As such, a comprehensive phylogeny and detailed morphological study including more samples from across the range of the genus is required to revise the subgeneric classification of the genus.

Notwithstanding the unclear sectional classification of the genus in Australia, the resolution of these three major clades in our phylogeny has interesting implications for the biogeographic evolution of the genus in Australia when taken in the context of previous work. Maurya *et al*. (2023), reconstructed the ancestral area for *Capparis* as Southeast Asia or India, and *C. anomala*, *C. sepiaria* (from a Philippines locality), and the clade of *C*. sect. *Busbeckea* were not monophyletic, suggesting at least three independent range expansions to Australia occurred in the radiation of this clade. In our analysis, the retrieval of the second clade of *C*. sect. *Monostichocalyx* species and its position suggests that this lineage emerged from a separate range expansion into Australia from India or Southeast Asia, but this needs to be verified in the context of a phylogeny with broader, global sampling. As we found *C. spinosa* subsp. *cordifolia* (which only occurs outside of Australia) to be sister to the *C. spinosa* subsp. *nummularia* clade, our phylogeny implies that there was at least one additional range expansion of *C. spinosa* to Australia. The origins of *C. spinosa* in Australia have long been debated, with Jacobs (1964) postulating that it was introduced to Australia by humans from Europe ‘somewhere on the NW coast’, but later authors considering it to be autochthonous (Hewson 1982; Fici 2003). Our results support an independent range expansion of *C. spinosa* into Australia, but determining its timing, mode and locality requires more comprehensive sampling of *C. spinosa* subspecies across its range, and a well-dated tree. In sum, our results indicate it is likely that there were at least five independent range expansions of *Capparis* into Australia, which resulted in different patterns of diversification and evolution.

We detected a WGD event at the crown node of *Capparis*. This is in line with karyological (Lysak 2018), transcriptomic (Mabry *et al*. 2020) and genomic (Wang *et al*. 2022) studies, which have inferred an ancient WGD event in the evolution of Capparaceae. Locating the exact position of this event requires additional data and sampling from across the family, but our analysis supports this event having occurred before the evolution of the most recent common ancestor of Australian *Capparis*.

In the clade comprising the *C.* sect. *Monostichocalyx* species *C. quiniflora*, *C. lasiantha*, *C*. sp. Bamaga (V.Scarth-Johnson 1048A), *C. sarmentosa*, and *C. batianoffii*, the signature of this ancient WGD event is still present, with moderate levels of allele divergence and heterozygosity, and some minor phylogenetic discordance in the topology of *C. lasiantha*, *C*. sp. Bamaga (V.Scarth-Johnson 1048A), *C. sarmentosa*, and *C. batianoffii*. Nevertheless, the consistency in the topology of the trees across analyses in this clade suggest that these species are divergent enough to reconstruct phylogenetic relationships in the face of this ancient WGD event when paralogs are carefully handled. This genetic divergence is supported by the fact that these species are relatively distinct and easier to distinguish from each other.

In the *C.* sect. *Capparis* clade, heterozygosity and allele divergence was low, suggesting that this clade may have undergone dysploidy or diploidisation in Australia (Nauheimer *et al*. 2021; Hendriks *et al*. 2023). However, the high allele divergence and heterozygosity of *C. spinosa* subsp. *nummularia* 6 and *C. spinosa* subsp. *cordifolia* indicates that these individuals are polyploids, and that polyploidy readily occurs in the species. The retrieval of three clades that are consistent with geographical isolation and morphology indicate that these clades represent lineages that are genetically distinct, even at a within-species level. The estimation of two alternative topologies for these clades in our species trees suggests a recent radiation characterised by incomplete lineage sorting. Nevertheless, our molecular evidence suggests that there are three, genetically divergent lineages in *C. spinosa* subsp. *nummularia* that correspond with both morphology and geography.

The evolution of these clades, whereby only a signal of ancient WGD persists amongst relatively divergent lineages, contrasts to the evolution of the *C.* sect. *Busbeckea* clade. A second WGD duplication event was inferred at the crown of this clade, which explains the extremely high allele divergence and heterozygosity of species in this clade. The exceptions to this are ‘*C*. sp. hairy-leaved form’, *C. thozetiana*, and *C. umbonata*, for which we had a lower coverage of data and therefore could probably not extract as many paralogs. We hypothesise that the polyploidisation event at the base of *C*. sect. *Busbeckea* resulted in rapid diversification of lineages, which along with the issue of paralogs in this clade, has resulted in the high lability in species relationships across trees, poor topological support, short branch lengths and phylogenetic discordance. This also explains the complex patterns of variation and the difficulty in differentiating species in this clade. We did not test whether this WGD event was the result of a hybridisation or autopolyploidisation event; given that our samples were restricted to Australia, we probably have an unbalanced tree and are potentially missing parental lineages, making our dataset unsuitable for differentiating between these causes. Therefore, the nature of the WGD event at the crown of *C*. sect. *Busbeckea* remains to be identified with a more complete dataset.

### Classification of Australian *Capparis*

The complex evolutionary history of *Capparis* in Australia has interesting implications for taxonomy. Ancient WGD events have likely caused rapid radiation and complex patterns of morphological variation, making taxonomic boundaries at the species level difficult to reconstruct using nuclear molecular data or morphology alone. Therefore, it is especially important to consider molecular data, geography, morphology, and ecology together to determine species boundaries in *Capparis*. We discuss the taxonomic status of each of the sections, phrase-named taxa, putative new taxa, and complex species for Australia below. A taxonomic treatment of new species, and identification key to all Australian *Capparis* taxa follows.

In the interests of taxonomic stability, we have chosen not to recircumscribe sections of *Capparis* for Australia. Although *C*. sect. *Monostichocalyx* is not monophyletic, we suggest that formal recircumscription of sections and identification of morphological synapomorphies should follow a molecular and morphological study with comprehensive sampling from across the genus’ range.

Our analyses have shown that both phrase-named taxa — *Capparis* sp. Bamaga (V.Scarth-Johnson 1048A) and *C*. sp. Coen (L.S.Smith 11862) — warrant recognition as species. *Capparis* sp. Bamaga (V.Scarth-Johnson 1048A) was retrieved as a distinct branch in the *C*. sect. *Monostichocalyx* clade, most closely related to *C. sarmentosa*, *C. batianoffii* and *C. lasiantha*. We herein describe it as *C. xylocarpa* W.E.Cooper. *Capparis* sp. Coen (L.S.Smith 11862) was found to belong to the *C*. sect. *Busbeckea* clade, and was consistently retrieved as sister to the rest of *C*. sect. *Busbeckea.* As such, we erect the name *C. megacarpa* W.E.Cooper for this species.

Both putative new species — ‘*C*. sp. hairy-leaved form’ and ‘*C*. sp. Wet Tropics vine’ — were placed in the *C*. sect. *Busbeckea* clade. Given the complex evolutionary history of the clade with a whole-genome duplication event and rapid radiation, it is perhaps unsurprising that the exact relationships of these taxa differed across species trees. However, the trees do indicate that they are as genetically distinct as other species in this clade. Moreover, they can be clearly distinguished on morphological and ecological character states, and we therefore propose that these entities merit recognition as distinct species. As such, we formally describe ‘*C*. sp. hairy-leaved form’ as *C. loxophleba* W.E.Cooper & Joyce, and ‘*C*. sp. Wet Tropics vine’ as *C. splendidissima* W.E.Cooper below.

*Capparis lucida* is always retrieved as a clade in the *C*. sect. *Busbeckea* clade, confirming the monophyly of this species. Although the relationship of *C. lucida* differs across species trees, it is always distinct from *C. arborea*, in contrast to the suggestion of Hewson (1982). We suggest that the morphological similarity between *C. lucida* and some *C. arborea* individuals is the consequence of the history of WGD and rapid radiation in the *C*. sect. *Busbeckea* clade.

*Capparis arborea* is not monophyletic in any of our species trees, with the northern individuals consistently retrieved separately from the southern individual. This was poorly supported, and the sister relationships of the *C. arborea* individuals also differed across trees due to the complex evolutionary history of sect. *Busbeckea*. *Capparis arborea* also displays complex and subtle morphological variation across its range in indumentum density, hair orientation, and leaf shape, causing confusion with similar species such as *C. lucida* in parts of its range. However, after careful examination of specimens from CNS, BRI and MEL and field observations, we have not been able to identify any clear morphological or ecological characters that consistently distinguish *C. arborea* across its range. Considering this, in combination with the unclear genetic evidence, we have decided to retain the current concept of *C. arborea*. Future genomic or transcriptomic studies with additional samples and power might resolve *C. arborea* into distinct genetic entities that may warrant taxonomic recognition; however, we cannot justify splitting this species based solely on our molecular evidence due to the history of nested whole-genome duplications and rapid radiation.

The two subspecies of *Capparis loranthifolia* are consistently recovered as non-monophyletic, suggesting that each subspecies is more closely related to other species than to each other. Therefore, in agreement with the suggestion of Hewson (1982) and distinct morphological characters, we raise C*. loranthifolia* var. *bancroftii* to species level as *C. bancroftii*.

Our analysis shows that *Capparis spinosa* subsp. *nummularia* is monophyletic, and distinct from the morphologically and geographically close *C*. *spinosa* subsp. *cordifolia*. As discussed above, it also indicates that there are three recently diverged but genetically distinct clades in Australia; the WA lineage, the inland NT+QLD lineage and the island E.QLD clade. Fici (2003) concluded that the morphological differences between Western Australian and Northern Territory individuals were environmentally driven, and that these forms do not warrant taxonomic recognition. However, this finding was heavily based on observation of seedling morphology, and given that our data shows that separation of these lineages is likely recent, it is expected that there is overlap in some morphological characters. This is the case in subpecies of *C. spinosa* in the Mediterranean region, which have overlapping morphological characters, and are known to hybridise in sympatry (Fici 2014; Gristina *et al*. 2014). Due to their morphological differences, geographical isolation and genetic distinctiveness, we argue that *C. spinosa* in Australia comprises three separate lineages, and we herein recircumscribe it into three subspecies. Subspecific rank is in keeping with the taxonomic concepts of the broader species worldwide, and also recognizes the recency of the divergence of these three taxa. The type of *C. spinosa* subsp. *nummularia* is from Western Australia, and as such the WA form of *C. spinosa* in Australia retains the subspecific epithet *C. spinosa* subsp. *nummularia*. For the NT+QLD form from the Northern Territory and inland Queensland we raise *C. nummularia* var. *minor* Domin to subspecific rank as *C. spinosa* subsp. *formicosa W.E.Cooper & Joyce*, and erect *C. spinosa* subsp. *insularis* W.E.Cooper & Joyce for the E. QLD form from the islands off northern Queensland. Further morphological and molecular study, with comprehensive sampling across the subspecific taxa and range of *C. spinosa* is needed to confirm the relationships of the Australian subspecies, and whether they have diversified in Australia following one or multiple range expansions. Such a study, in the context of a broader phylogenetic study of the genus, would also help to inform whether some *C. spinosa* subspecies and varieties of subspecies should be elevated to specific level.

### Taxonomic treatment

*Type*: *Capparis spinosa* L.

***Capparis*** L., *Sp. Pl*. 1: 503 (1753).

In Australia: Monoecious (or dioecious only in *C. anomala*) leafy shrubs (with the exception of *C. anomala* which is often leafless), trees, scramblers, vines or lianas. Indumentum mostly present, hairs simple or rarely stellate or T-shaped. Stipular spines paired, persistent, straight or recurved, often not apparent on new growth. Leaves simple, alternate and distichous. Inflorescence axillary, terminal or ramiflorous, a solitary flower, serial rows, umbellate, racemose or paniculate. Flowers nocturnal in species within sections *Capparis* and *Busbeckea*, pedicellate, zygomorphic, 4-merous; sepals in 2 whorls, outer pair free or connate, inner pair free. *Petals* 4, fragile; stamens few to *∼*200; gynophore present, ovary 1-locular, ovules few– many, stigma sessile; fruit a balausta (hard and indehiscent) or a capsule on a gynophore attached to a pedicel, seeds few–many, embedded in fleshy or dry pulp, obliquely reniform. Germination epigeal.

### Etymology

*Capparis* is from Kapar, the Arabic name for *Capparis spinosa*.

Capparis L. sect. *Capparis*

*Type species*: *Capparis spinosa* L.

*Capparis* sect. *Eucapparis* DC., *Prodr*. 1: 246 (1824), *nom. inval. Capparis* sect. *Eucapparis* Benth., *Fl*. *Austral*. 1: 93-95 (1863), *nom. inval*.

**Capparis spinosa L.**, *Sp. Pl.* 1: 503 (1753)

*Type citation*: “Habitat in Europae australis arenosis, ruderalis”. *Lectotype*: designated by Burtt & Lewis in *Kew Bull*. 4: 299 (1949): Herb. Clifford: 203 (BM BM000628727).

In Australia: *Shrub* to 2 m, erect, spreading, prostrate or decumbent; stems may be zigzag. *Indumentum* on stems and leaves lanulose, glabrescent, white. *Stipular spines* mostly on older stems, rarely on flowering stems, straight or slightly recurved, sometimes erect, 0.5–5 mm long, yellow or orange. *Leaves* simple, alternate, entire; petioles 4–18 mm long, grooved; lamina orbicular, obovate, broadly elliptical or ovate, thinly succulent; apex mucronate; venation brochidodromous; secondary veins 3–7 pairs, tertiary veins reticulate. *Inflorescence* axillary, a solitary flower. *Flowers* often fragrant; zygomorphic. *C alyx* 4-merous, 2-whorled, posterior sepal galeate, anterior sepal boat-shaped; apex mucronulate; inner sepals 2, boat-shaped. *Petals* 4, white, ageing to pink or purple, obovate or flabellate. *Stamens* numerous, up to ∼ 120; anthers grey; *ovary* ellipsoidal or spindle-shaped, 6-ribbed. *Fruit* pedicel and gynophore usually loosely spiralled; capsule fleshy, 5–7-ribbed or 5–7-cornered, dehisces tardily and at random either along ribs or in between the corners or ribs; seeds numerous, asymmetrically reniform or snail-shape, suspended in aril.

*Capparis spinosa* subsp*. nummularia* (DC.) Fici

*Capparis nummularia* DC., *Prodr. 1*: *246* (1824). *Type citation*: “in Novae-Hollandiae insulus sterilibus (v.s. sine fl. et fr.)”. *Type:* AUSTRALIA. WESTERN AUSTRALIA. Bernier & Dorre Is. (Iles Stériles), *s. dat., leg. ign., s.n.* (holo: G-DC G00207304; iso: L L0035357, P P00337390). *Capparis nummularia* var. *typica* Domin (1925: 686). *nom. inval*.

### Illustrations

Wheeler *et al*. (1992, p. 73:F). Milson J, (2000: 38–39).

*Stipular spines* on flowering stems as well as on older growth, slightly recurved or straight and flanged at base, 0.5–3 mm long, yellow or orange. *Leaves*: petioles 6–18 mm long, lanulose, glabrescent; lamina orbicular or obovate, rarely ovate, 12–43 mm long and 10–47 mm wide, glabrescent adaxially and abaxially, glaucous; base cuneate or rounded, rarely shallowly cordate; apex mostly shallowly emarginate, rarely rounded, mucronate, mucro mostly recurved or underdeveloped as a short wart, usually emerging at right angles to leaf and only visible abaxially, 0.25–1.25 mm long; primary vein flush adaxially and raised abaxially; secondary veins 3–7 pairs diverging at 50–90° to the primary vein, slightly raised adaxially and distinctly raised abaxially. *Inflorescence*: pedicel 15–60 mm long, thinly lanulose. *Flowers*: bud apex acuminate, obtuse or bluntly acute and mucronulate, mucro up to 0.25 mm long on sepals and soon deciduous, sepals ribbed or not ribbed, lanulose, soon glabrescent. *Calyx* green, maybe be purplish on margins; posterior sepal 12–30 mm long and 7–30 mm wide; outer anterior sepal 12–20 mm long and 6–15 mm wide, glabrous, apex may be cuspidate with a mucro up to 0.3 mm long; inner sepals 12–20 mm long and 6–15 mm wide, glabrescent abaxially, adaxially lanulose, glabrescent with a few sparse hairs remaining at base. *Petals*: posterior pair 20–30 mm long and ∼25 mm wide; abaxially with sparse hairs and adaxially glabrous (Fig. 4). *Stamens* numerous ∼ 80, filaments *∼*16–50 mm long, white aging to pink; anthers 2*–*2.5 mm long. *Gynophore* 35–45 mm long, minutely and thinly lanulose in lower half, glabrescent. *Ovary* 4.5–5.5 mm long and 2–2.25 mm wide, glabrous or with sparse hairs. *Fruit*: pedicel 45–60 mm long, glabrous; gynophore 45–50 mm long, thinly lanulose, glabrescent; capsule ovoid, orange or yellow, 5–7-corned (immature fruit ribbed, ripe fruit not ribbed), 45–58 x 20–28 mm, glabrescent; seed diameter 3–4.25 mm long, dark brown, suspended in bright red aril (or yellow) (Fig. 4).

**Fig. 3.**
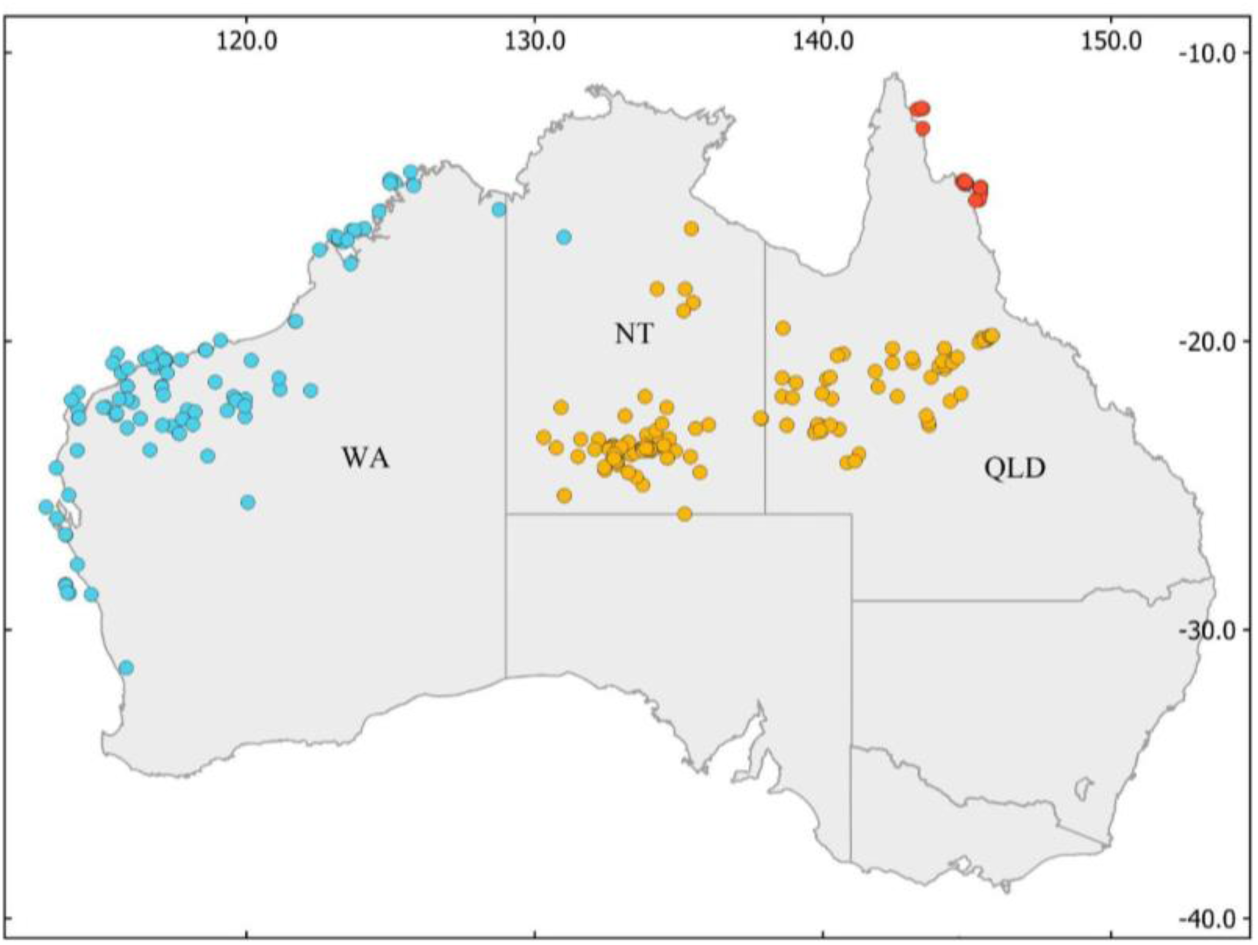
Capparis spinosa distribution in Australia: 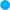*Capparis spinosa* subsp. *nummularia*; 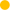 *Capparis spinosa* subsp. *formicosa*; 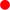 *Capparis spinosa* subsp. *Insularis*.

**Fig. 4.**
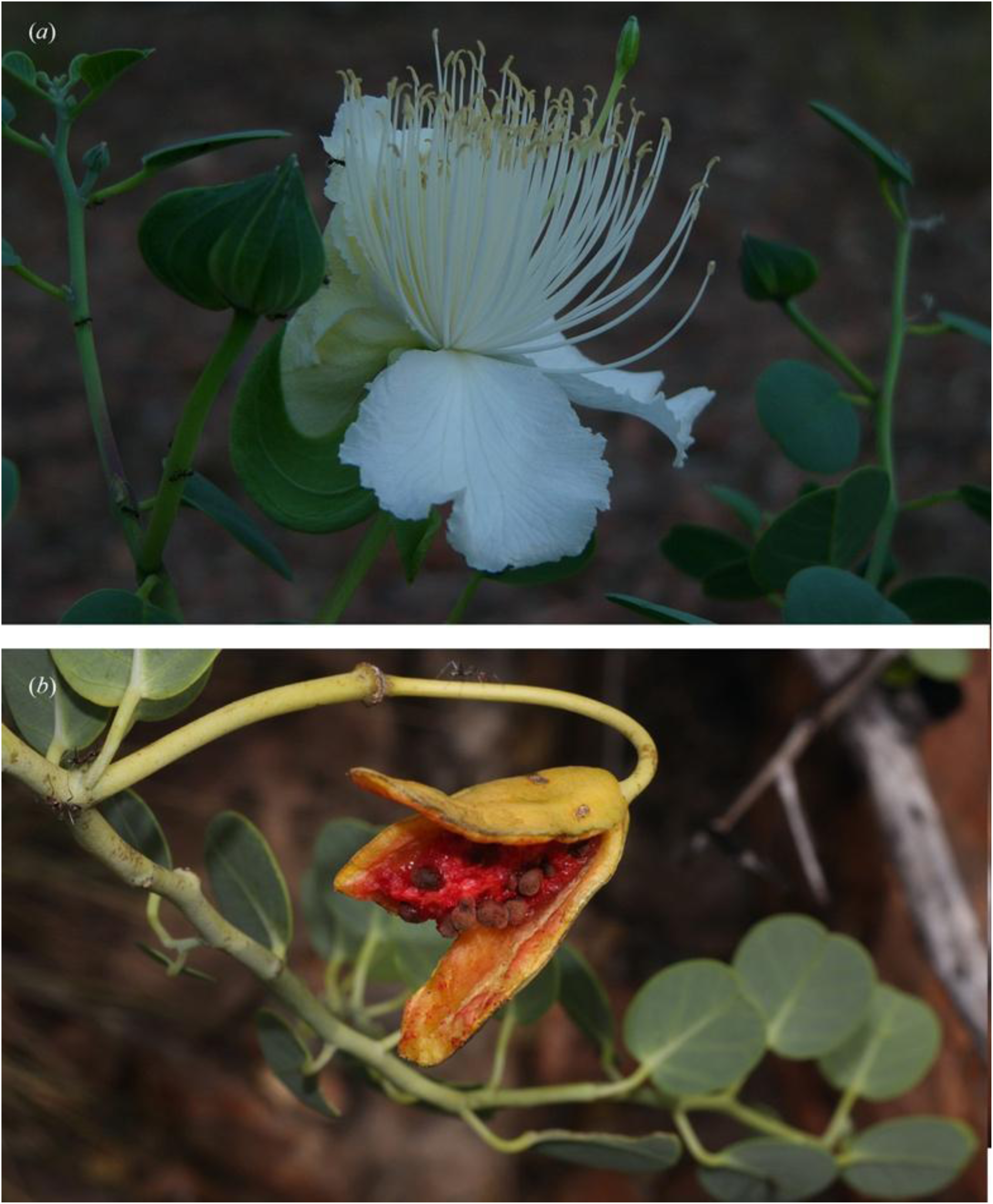
*Capparis spinosa* subsp. *nummularia*: (*a*) Flower in anthesis showing galeate anterior sepal with remaining sepals boat-shaped. (*b*) Dehisced fruit exposing red aril and brown seeds. Photographs: (*a*) T. Start, (*b*) R. Barrett.

### Distribution

*Capparis spinosa* subsp. *nummularia* occurs in coastal areas of Western Australia, north from Geraldton to Fenelon Island on the Kimberley coast, as well as inland in the greater Pilbara region and also with one record from Victoria River Downs in the Northern Territory (Fig. 3).

### Habitat and ecology

*Capparis spinosa* subsp. *nummularia* occurs in various habitats from stony hillsides, sand, pavement limestone, rocky escarpment, heath, *Triodia* grassland, shrubland and open woodland.

### Phenology

Flowers have been recorded from January to November and fruit recorded in April and June.

### Etymology

The specific epithet *nummularia* is from the Latin *nummus* (coin) in reference to the round or coin-shaped leaves.

### Specimens examined

WESTERN AUSTRALIA. Banks of Ashburton River at Balkra between Globhill and Minderoo, 7 Oct. 1905, *A.Morrison s.n*. (BRI, CANB, PERTH); Port Walcott W.A., *s.dat.*, *F.Mueller s.n.* (NSW); Millstream, 90 miles NW of Wittenoom, Sep. 1969, *M.I.H.Brooker 2083* (NSW, PERTH): Port Headland, Apr. 1905, *W.F.Fitzgerald s.n.* (NSW); On mine road south of Paraburdoo, Jan. 1979, *P.A.Lascelles s.n.* (CANB); Ca. 9.2 km SE of Wyloo [Homestead] turnoff, 2 May 1977, *Hj.Eichler 22575* (CANB, DNA, PERTH); Dampier, Fortesque district, Apr. 1979, *Z.Smithson s.n.* (CBG); Dirk Hartog Island, Jan. 1822, *A.Cunningham 234* (BM, K); Roy Hill Station, 8 May 1958, *N.T.Burbidge 6024* (CANB); Monte Bello Is., 12 Nov. 1953, *P.Hitchin s.n.* (CANB); Road from North West Coast Highway to Exmouth, at Boundary Creek which is W of Giralia HS turnoff, 8 June 2002, *A.V.Slee 4465* (CANB); Long Island, Buccaneer Archipelago, 13 June 1982, *A.J.M.Hopkins BA 0106* (PERTH); Port Sampson, 18 Oct. 1941, *C.A.Gardner 6315* (PERTH); 44 km SSW of Nullagine, 31 Mar. 1984, *K.R.Newbey 10091* (PERTH); Mundiwindi septentrionalem versus pr. 807.5 mile peg [N of Mundiwindi, near 807.5 mile peg], 17 Oct. 1963, *F.Lullfitz 2686* (PERTH); 2 miles from Ashburton River, 20 Oct. 1970, *J.R.Cannon 115* (PERTH); Barrow Island Nature Reserve, John Wayne Country, 7.8 km SSE of The Castle, 11.3 km WNW of Town Point, 13 Nov. 1991, *S. van Leeuwen 1069* (PERTH); Bigge Island, 1 June 1972, *N.G.Marchant 72/69* (PERTH); Near Roebourne, Apr. 1901, *E.Pritzel 284* (L, NSW); Cossack, N of Roeburn, Apr. 1901*, F.L.E.Diels & E.Pritzel* (PERTH); Legendre Island, Dampier Archipelago, 9 June 1962, *R.D.Royce 7245* (PERTH); Crusher Vine Thicket, Mitchell Plateau, N Kimberley, Sep. 1990, *R.A.McKenzie s.n.* (PERTH); Little Pigeon Island, Wallabi Group, Abrolhos Islands, 20 Mar. 2003, *V.Casey 1* (PERTH); North Sandy Island, Oct. 1949, *D.L.Serventy s.n.* (PERTH). NORTHERN TERRITORY. Walsh Creek, Victoria River Downs, 24 Nov. 1968, C.S.Robinson 49(DNA).

***Capparis spinosa*** subsp*. **formicosa*** W.E.Cooper & Joyce, *subsp.. nov*.

T*ype:* AUSTRALIA. QUEENSLAND. Burke District: Hughenden, 20 Nov. 2020, *Cooper 2698 & Ford* (holo: CNS 155749 [2 sheets + spirit], iso: 7 sheets to be distributed to BRI, CANB, PERTH, L, M, MO, PR).

*Capparis nummularia* var. *minor* Domin, *Biblioth. Bot.* 22(89): 686 (1926). *Type citation:*

”Queensland: am Flinders River bei Hughenden (DOMIN II. 1910).”.

*Capparis spinosa* var. *nummularia* F.M.Bailey, Syn. Queensland Fl.: 15 (1883). Type: [not cited].

### Illustration

Latz (1995), p. 141.

### Diagnosis

*Capparis spinosa* subsp*. formicosa* is similar to *Capparis spinosa* subsp*. nummularia* but differs in the leaves mostly ovate or elliptical, rarely orbicular or obovate (*v*. orbicular or obovate, rarely ovate); leaf apex mucro 1–3 mm long (*v*. 0.25–1.25 mm long); sepals tomentose to glabrescent (*v*. glabrous); petals 35–38 mm long (*v*. 20–30 mm long); fruit pedicel 35–45 mm long (*v*. 45–60 mm long); fruit ribbed (v. cornered, not ribbed); aril orange (*v*. red or orange).

### Description

*Shrub,* erect, to 2 m; bark grey with pale longitudinal lenticels. *Stipular* spines present on sterile stems and on flowering stems, recurved, 0.5–5 mm long, orange becoming grey on old wood. *Leaves*: petioles 4.5–11 mm long, glabrescent; lamina mostly ovate or elliptic, rarely obovate or orbicular, 10–65 mm long and 8–42 mm wide, abaxially with a few sparse minute white hairs, glabrescent, adaxially glabrous, glaucous; base truncate, cuneate or rounded; apex retuse, shortly acute, rounded, emarginate or acute, mucronate, mucro a recurved orange thorn 0.3–3 mm long and extending on the same plane as the midrib and visible from above; primary vein raised on abaxial surface and ± flush adaxially; secondary veins 4–6 pairs, flush on both sides or slightly raised abaxially, angle of divergence from primary vein 20–60°. *Inflorescence*: pedicel 20–50 mm long, lanulose, glabrescent. *Flowers*: bud apex shortly acuminate or shortly cuspidate, sepals in bud have 5 distinct longitudinal veins which are raised as 5 ribs which become indistinct in open flowers and dried specimens, pink to maroon-coloured nectiferous glands are exposed on bud expansion with one on each side at base of outer sepals and a pair in the centre at base of inner sepals (these glands are usually not conspicuous in dried specimens). *Calyx* purplish to green; posterior sepal 18–36 mm long and 18–31 mm wide; anterior sepals 14–26 mm long, 5–8 mm wide, all have an apex shortly acuminate or mucronulate with a recurved yellow caducous thorn 0.15–1 mm long, abaxially lanulose in bud, glabrescent, adaxially glabrous. *Petals*: posterior pair 35–38 mm long and 25–30 mm wide, sparsely pubescent; anterior pair rhomboid, elliptical or ovate-elliptical, 14–35 mm long and 8–20 mm wide, glabrous; gland at base of proximal petals, diameter *c*. 4.5 mm. *Stamens* 100–120, filaments 28–45 mm long, white, anthers 2–3.5 mm long. *Gynophore* 10–60 mm long, sometimes much shorter than stamens, sparsely pilose towards base, glabrescent. *Ovary* 3.5–6 mm long and 2–2.5 mm wide, 6–8-ribbed, sparsely hairy or glabrous (Fig. 5). *Fruit*: pedicel 35–45 mm long, with sparse minute hairs or glabrous; gynophore often spiralled, up to 47 mm long, glabrous; capsule ovoid, up to 58 mm long and 29 mm wide, 6–8-ribbed, apricot-coloured or orange (often with green ribs) with dark reddish flush towards base especially on ribs, viscid becoming glabrous; seed diameter *∼* 4 mm, dark brown, suspended in orange aril (Fig. 5). *Caper Bush, Flinders Rose, Wild Passionfruit, Split Jack, Wild Nipan, Wild Guava, Ant Bush*

**Fig. 5.**
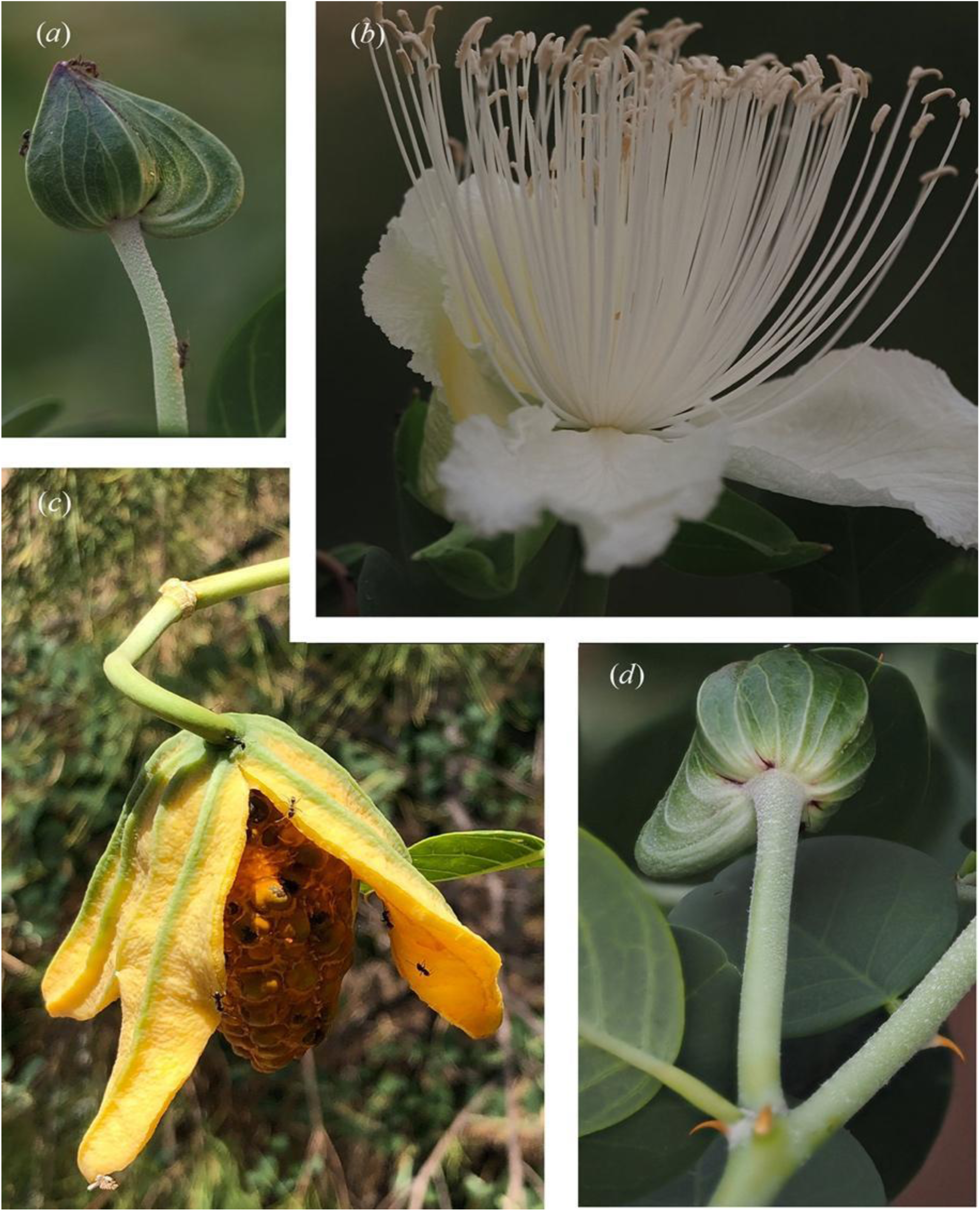
*Capparis spinosa* subsp. *formicosa*: (*a*) Bud showing ribbed sepals, mucronulate apex and lanulose pedicel. *Cooper 2965.* (*b*) Flower at anthesis. *Cooper 2965.* (*c*) Dehisced fruit showing orange aril and brown seeds. *Cooper 2735.* (*d*) Stem showing stipular spines, bud with ribbed sepals, dark red glands at sepal bases and lanulose pedicel. *Cooper 2965.* Photographs: W. Cooper.

### Distribution

*Capparis spinosa* subsp. *formicosa* is known in the Northern Territory, north from the border with South Australia to Mallapunyah in Arnhemland. In Queensland it occurs north from the Diamantina Lakes and east towards the western fall of the Great Dividing Range (Fig. 3).

### Habitat and ecology

*Capparis spinosa* subsp. *formicosa* occurs on various soils in low woodland, savannah and grasslands including Mitchell Grass plains, sandy desert, exposed or disturbed and degraded areas such as roadsides, rocky ridges and gorges, eroded gullies, heavily grazed grassland and seasonal watercourses.

### Phenology

Flowers have been recorded from every month and fruit from November to July.

### Etymology

The specific epithet *formicosa* is derived from the Latin meaning full of ants; referring to the ever present ants on all parts of these plants.

### Notes

The Type of *Capparis nummularia* var. *minor* Domin was collected by Karel Domin, Feb. 1910, at Flinders River near Hughenden, Queensland. That specimen is apparently lost, and duplicates have not been located (pers. comm. Ota Sida, National Museum Prague).

Bailey (1883:15) in his *Synopsis of the Queensland Flora* did not cite a type for *Capparis spinosa* var. *nummularia* F.M.Bailey, and made no reference to a basionym, and therefore the name has been treated as a *nom. nov.*, even though it is likely to have been based on *C. nummularia* DC. He used abbreviations “*I. Int.*” to indicate that the plants occurred in Inland Intratropical regions and given the geographic scope of the book we can surmise that the concept was applied to specimens that we are herein treating as *Capparis spinosa* subsp. *formicosa*.

*Capparis spinosa* subsp. *formicosa* flowers begin senescing between early and late morning, withering and turning pink to magenta. At sunrise, flowers (especially the anthers) are visited by exotic honey bees (*Apis* spp.). *Capparis spinosa* subsp. *formicosa* is often heavily grazed by cattle and occasionally by horses.

### Specimens examined

AUSTRALIA. QUEENSLAND. 16km by road west of Boulia on Donoghue Hwy, 27 Mar. 2007, *M.B.Thomas 3148 & G.P.Turpin* (BRI, DNA); Astrebla Downs National Park, *c.* 140 km from Bedourie, 10 June 2005, *M.Mostert MM82* (BRI); 11km N of Phosphate Hill mine, 8km SE of the Monument township, 1 May 1985, *V.J.Neldner 2058 & T.D.Stanley* (BRI); Toorak Field Station, S of Julia Creek, 6 May 1956, *N.T.Burbidge 5345* (CANB); Entrance to Eldorado, Winton to Hughenden Road, 3 Mar. 2021, W.E.*Cooper 2748 & A.J.Ford* (CNS, BRI, CANB, NSW); Roadside, northern approach to Hughenden, 18 Dec. 2020*, W.E.Cooper 2735 & B.Barnett* (CNS); Torver Valley Road, Hughenden, 7 May 2023, *W.E.Cooper 2965* & *B.Barnett* (CNS, BRI, CANB); Toomba, 27 July 2020, *W.E.Cooper 2643, R.Jensen & F.A.Zich* (CNS, BRI, CANB); Boulia, *s.dat.*, F.M.*Bailey s.n.* (NSW); *Ca.* 20 miles SE of Kynuna, Landsborough Highway between Longreach and Cloncurry, 16 Aug. 1967, *B.G.Briggs 1177* (NSW); NORTHERN TERRITORY. Palm Creek, Finke Gorge, 16 miles NW of Hermannsburg Mission N.T., 22 Mar. 1953, *R.A.Perry 3508* (BRI, CANB, DNA, PERTH); Emily Gap, about 6 miles east of Alice Springs, 22 Jan. 1950, *S.L.Everist 4154* (CANB, BRI); 22 km W of Numery Homestead, 14 Sep. 1993, *P. Latz 13343* (NT).

The collection *Tepper 15* (MEL 590852), is a good match for *Capparis spinosa* subsp. *formicosa*. However, two collection sites are recorded on this herbarium sheet: Lake Eyre country and Kings Sound, WA, which is potentially confusing. Kings Sound is an unlikely collection site with morphology being typical of plants occurring in the Northern Territory and inland areas of Queensland. Lake Eyre is at least 300 km from the most southerly reliable record in central Australia: *Latz 13343* (22 km W of Numery Homestead, NT) near the NT-SA border. The apparently tireless plant collector, Johann Gottlieb Otto Tepper (1841–1923), collected in regions of South Australia: Yorke Peninsula, Barossa Valley, Mount Lofty Ranges, Murray Mallee and Kangaroo Island but is not recorded as having visited Lake Eyre. It could therefore be hypothesized that the collection MEL 590852 was collected from within the loose location referred to as Lake Eyre country which might have applied broadly to the arid center.

***Capparis spinosa*** subsp*. **insularis*** W.E.Cooper & Joyce, *subsp. nov*.

*Type*: AUSTRALIA. QUEENSAND. Cook District: Restoration Rock, 1st Dec. 2018, *W.E. Cooper 2577, J. Pritchard & B. Venables* (holo: CNS 152674 [3 sheets + spirit], 8 sheets to be distributed to BRI, CANB, NSW, L, K, MO, P).

### Diagnosis

*Capparis spinosa* subsp*. insularis* is similar to *Capparis spinosa* subsp. *nummularia* but differs in the leaf apices mostly shortly acute (v. mostly retuse); gynophore 65 mm long (*v*. 40–50 mm long); fruit pedicels 60 mm long (*v*. 30–60 mm long); fruits distinctly 5-ribbed (*v*. 5-cornered); aril orange (*v*. red or orange).

*Capparis spinosa* subsp*. insularis* is similar to *Capparis spinosa* subsp. *cordifolia* (Lam.) Fici but differs in having stipular spines usually present (*v*. rarely present); leaves ± orbicular, ovate-orbicular or rarely broadly elliptical (*v*. mostly ovate, elliptical or rarely sub-orbicular); leaf apex mucronulate with a recurved spine (*v*. minutely mucronulate, the mucro only a small protruberance or wart-like and not a recurved spine); flower bud apex shortly acuminate or shortly mucronate (*v*. obtuse); ovary ellipsoid (*v.* narrowly obovoid); fruit obovoid or broadly ellipsoid (*v*. oblong, narrowly ellipsoid or narrowly obovoid).

*Stipular spines* mostly present on sterile stems where they are up to 3.5 mm long, straight or slightly recurved, orange. *Leaves*: petioles 4–13 mm long, lanulose, glabrescent; lamina ± orbicular, ovate-orbicular or rarely broadly elliptical, 10–42 mm long and 7–45 mm wide, glabrescent abaxially and adaxially or with few sparse hairs mostly near base abaxially, glaucous; base cuneate, cordate, rounded, attenuate or truncate; apex mostly shortly acute, rarely rounded or retuse, mucronate, mucro a recurved orange thorn (rarely wart-like) 0.25– 0.75 mm long and attached at right angles to the lamina and usually only visible abaxially; primary vein slightly raised abaxially and adaxially; lateral veins 5–7 at 50–70°, raised on both sides. *Inflorescence*: pedicel 28–65 mm long, lanulose, glabrescent or with some hairs persisting near base; flower bud apex shortly acuminate or shortly mucronate, lanulose abaxially, glabrescent, maroon-coloured nectiferous glands exposed on bud expansion with one on each side at base of outer sepals and a pair in the centre at base of inner sepals (these glands are usually not conspicuous in dried specimens), buds with 5 raised longitudinal veins becoming indistinct in open flowers and dried specimens. *Calyx*: posterior sepal 20–32 mm long and 20–31 mm wide, glabrescent; anterior sepals 23–26 mm long, 8–12 mm wide, glabrescent, apex shortly acuminate or shortly mucronate with a recurved yellow caducous thorn, purplish to green. *Petals*: posterior pair oblique, obovate or rhomboid, 27–38 mm long and 10–30 mm wide; numerous; filaments *∼* 45 mm long, white; anthers ∼3 mm long. *Gynophore* ∼35 mm long, sometimes with a few sparse hairs in lower half. *Ovary* ellipsoid, 5–6 mm long and ∼2.5 mm wide, 6-ribbed, sometimes with sparse minute hairs. *Fruit*: pedicel *∼* 60 mm long; gynophore *∼* 65 mm long with sparse minute hairs along stem (not at base); capsule obovoid or broadly ellipsoid, 36–40 mm long and 15–24 mm wide, ribbed, orange, often with dark reddish flush towards base especially on ribs. *Seeds ∼* 2.5 mm long and ∼2.8 mm wide, dark brown, suspended in orange aril (Fig. 6).

**Fig. 6.**
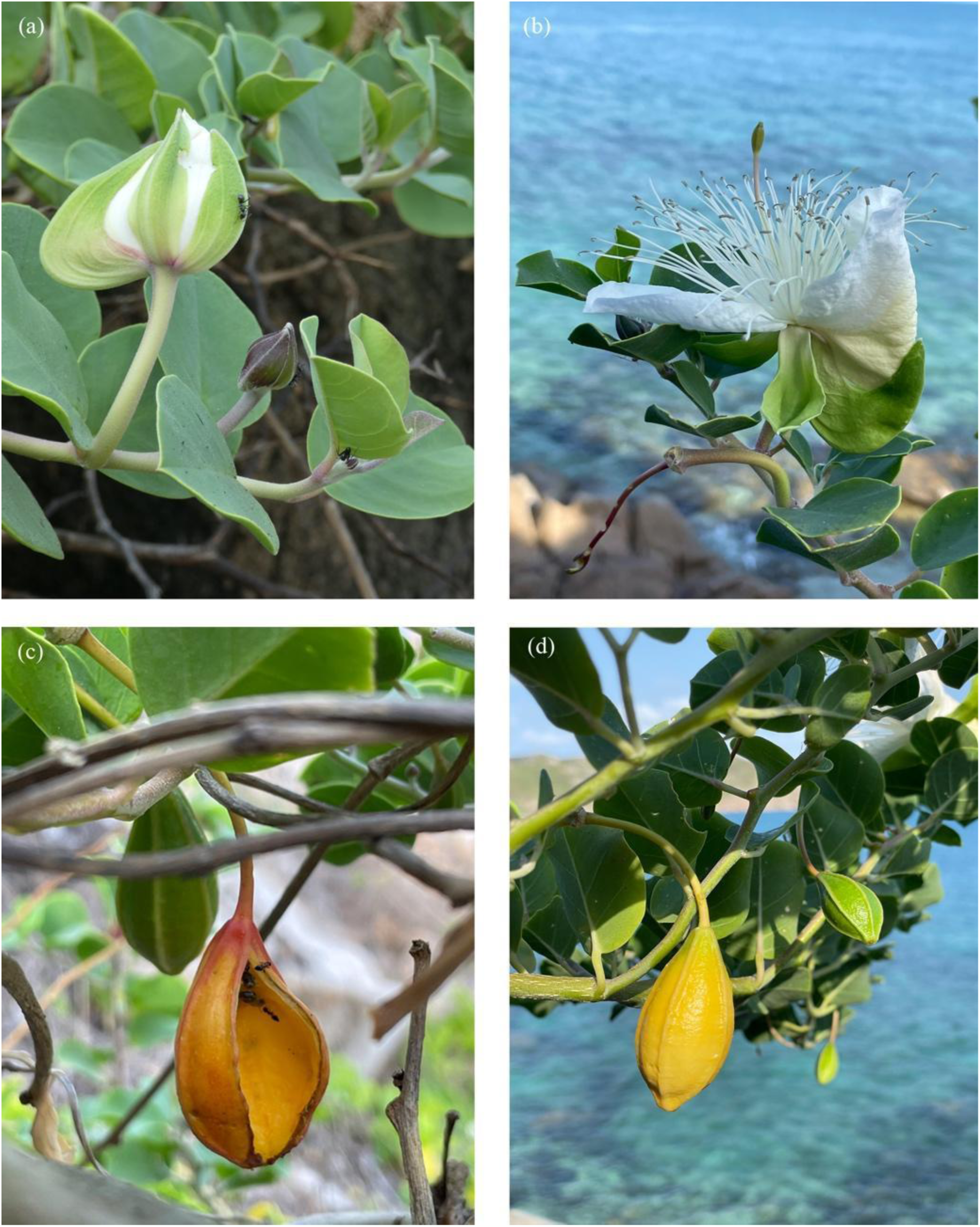
*Capparis spinosa* subsp. *insularis*: (*a*) Buds showing ribbed sepals and lanulose pedicels and sepals. *Cooper 2577*. (*b*) Flower at anthesis with galeate anterior sepal and boat-shaped inner sepal. (*c*) Dehisced ripe fruit. (*d*) Unripe indehisced fruit. *Cooper 3062*. Photographs: (*a*) W. Cooper; (b, c & d) J. De Raedt.

**Fig.7.**
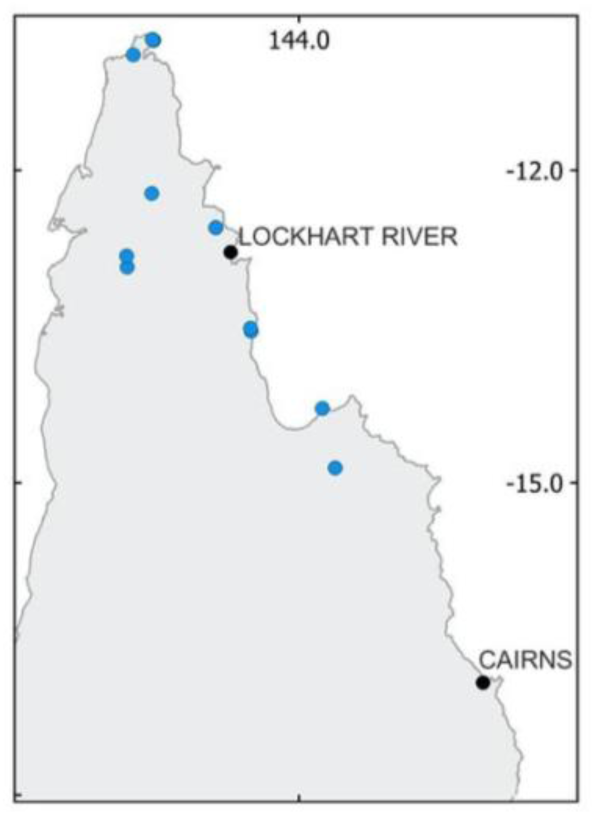
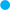 *Capparis xylocarpa* distribution.

### Distribution

*Capparis spinosa* subsp*. insularis* occurs on offshore islands in Queensland north from the Lizard Island group to the Home Island group (Fig. 3).

### Habitat and ecology

*Capparis spinosa* subsp*. insularis* is only known from continental islands and coral cays on various rock substrates including coraline sand, granite boulder fields and limestone and has not been recorded from the mainland. It grows commonly on exposed rock but also amongst herb and grassland as well as *Casuarina* woodland.

### Phenology

Flowers have been recorded for April, May, August, September, October, November and December. Fruit has been recorded for March, April, May and December.

### Etymology

The specific epithet is derived from the Latin *insularis* (pertaining to islands).

### Notes

At Restoration Rock, the Coastal Snake-eyed Skink *Cryptoblepharus litoralis* was observed eating the arils and seeds in ripe dehisced fruit.

### Specimens examined

AUSTRALIA. QUEENSLAND. Rocky Island, Lizard Island group, 31 May 2023, *W.E.Cooper 2984A, S.J.Worboys & I.A.Zorn* (CNS); Casuarina Beach, Lizard Island, 28 Sep. 1983, *Batianoff 10105* (BISH, BRI, DNA, MO); Lizard Island, 1 Aug. 1987, *N.Fisher 106* (CBG); Lizard Island, 24 Dec. 1974, *R.L.Specht LI353 & A.Specht* (BRI); Three Isles, 12 Sep. 1973, *D.R*.*Stoddart 4501* (BRI, L); Three Isles sand cay, 20 Nov. 2001, *B.M.Waterhouse BMW6324* (BRI, CANB); SW of Rocky Island, near South Direction Island, NE of Cape Flattery, Dec. 1989, *T.Walker s.n.* (BRI); Two Isles, 21 Sep. 1973, *D.R.Stoddart 4642* (BRI); Houghton Island, E of Cape Melville NP, 15 Nov. 1973, *I.R.Price 900059* (JCT); Howick Island, 22 Oct. 1973, *D.R.Stoddart 4847* (BRI); Howick Island, Flinders Group of Islands, 13 Dec. 1991, *G.N.Batianoff 911210A* (BRI); Sinclair Island, 1 June 2023, *W.E.Cooper 3016, S.J.Worboys & I.Zorn* (CNS); Sinclair Island, 8 Aug. 1973, *D.R.Stoddart 4188* (BRI, L); Coquet Island, Howick Group, 4 March 1983, M.*Godwin C2522* (BRI, CNS); Ingram Island, 27 July 1973, *D.R.Stoddart 4074* (BRI); Watson Island, 2 Aug. 1973, *D.R.Stoddart 4101* (BRI, L); Great Barrier Reef, Restoration Rock, near Cape Weymouth, Portland Roads, 24 July 1969, *Heatwole s.n.* (BRI); Restoration Rock, east of Cape Weymouth, 17 Oct. 2023, *W.E.Cooper 3062, J.Clarke, M.Paskins & J.De Raedt* (CNS); Perry Island, 10 Feb. 1991, *M.Card PER7* (BRI); Sir Charles Hardy Islands*, c*. 26 km ENE of Cape Grenville, 29 Nov. 1987, *J.R.Clarkson 7417* (BRI, CNS, MBA); Memey (Mimi) Island, 22km S of Masig (Yorke) Island, central Torres Strait, 26 Nov. 2016, *D.G.Fell DGFMM09, D.Baragud & E.Nai* (BRI).

*Capparis* sect. *Monostichocalyx* Radlk.

*Type species: Capparis micracantha* DC.

***Capparis xylocarpa*** W.E.Cooper, *sp. nov*.

*Type*: AUSTRALIA. QUEENSLAND. Cook District: Roadside near Somerset, 1 Sep. 2019, *W.E. Cooper 2585 & T. Hawkes* (holo: CNS 151465 [2 sheets + spirit], iso: 9 sheets to be distributed to BRI (2 sheets), CANB, DNA, K, L, MEL, MO, S).

*Capparis* sp. Bamaga (V. Scarth-Johnson 1048A) Qld Herbarium

*Capparis* sp. (Bamaga V. Scarth-Johnson 1048A) Jessup in Henderson (1997: 53), Jessup in Henderson (2002: 42), CHAH 2006, Jessup in Bostock & Holland (2007: 42), Guymer in Bostock & Holland (2010: 37), Guymer in Bostock & Holland (2013), Guymer in Bostock & Holland (2016)

*Capparis* sp. (Bamaga), Cooper & Cooper 2004: 110

### Illustrations

Cooper & Cooper (2004: 110)

### Diagnosis

*Capparis **xylocarpa*** is similar to *Capparis lasiantha* R.Br. ex DC. but differs in having stipular thorns 2–3 mm long (*v*. 3–4 mm long); petioles 3–6 mm long and 0.6–0.7 mm thick (*v*. 1.5–3 [rarely to 5] mm long and 1–1.7 thick); leaf shape ovate, broadly elliptical, subrhombic or broadly lanceolate (*v*. narrowly oblong, narrowly ovate or narrowly elliptical); leaves bright green, membranous, thinly-puberulent (*v*. yellowish, rusty or olive coloured, leathery, rusty velvety-puberulous); leaf apex mostly retuse (*v*. rarely retuse); reticulate veins distinct and intermittently swollen and blister-like (v. not visible); sepals 4.55 mm long (*v*. 6–12 mm long); sepals abaxially sparsely sericeous (*v*. tomentose). etals 8–9 mm long (*v*. 10–16 mm long). *Petals* abaxially sparsely sericeous (*v*. tomentose); stamens 16–18 (*v*. 20–22); fruit indehiscent (*v*. tardily dehiscent); fruit diameter 33–37.5 (*v*. 15–20 mm).

### Description

*Liana* to *∼*3 m, scrambling and sometimes decumbent, stem diameter up to *∼*4 cm, bark grey with numerous vertical fissures. Indumentum densely golden to rusty tomentose. *Stipular spines* recurved, to 2–3 mm long, orange to reddish-brown. *Leaves* discolorous; petioles 3–6 mm long, clothed in appressed hairs; lamina discolourous, ovate, broadly elliptical, subrhombic or broadly lanceolate, 20–55 mm long and 10–28 mm wide, membranous, both sides thinly puberulent with hairs becoming scattered, mostly with a tuft of hairs at apex abaxially growing from a wart; base cordate or rounded; apex tapering to a narrowly obtuse or retuse tip, entire; venation brochidodromous with the exception of the basal pair of veins, primary vein flush or slightly raised; secondary veins 4–6 at *∼*50°, slightly raised on both sides; reticulate veins intermittently swollen and blister-like. *Inflorescence* axillary, a solitary flower or rarely a serial row of 2 or a terminal raceme; pedicels 10–16 mm long, tomentose. *Flowers* without fragrance. *Calyx* slightly zygomorphic; sepals 4 in two whorls, cymbiform, 4.55 mm long and 4 mm wide, abaxially sparsely sericeous (sparser on inner pair), adaxially glabrous, green. *Petals* 4, oblong, recurved, *8*9 mm long and 4 mm wide, abaxially sparsely sericeous, adaxially glabrous, white. *Stamens* 16–18, filaments 18–20 mm long, white, anthers 2.5–3 mm long, grey. *Gynophore* 21–22 mm long, glabrous. *Ovary* narrowly ovoid, *∼*2.5 mm long and wide, glabrous; ovules numerous; stigma sessile, diameter *∼*0.35 mm; placentas 4. *Fruit*: pedicel 12–16 mm long, tomentose; gynophore 16–23 mm long, glabrous; balausta (hard and indehiscent) ellipsoid, 62– 80 mm long and 33–37.5 mm wide, warty, green; *seeds* several, diameter *∼*6 mm, dark brown, suspended in hard cream-coloured mesocarp. Germination epigeal. (Fig. 7)

### Distribution

*Capparis xylocarpa* is recorded intermittently on Cape York Peninsula between Lakefield National Park and Somerset near the tip of Cape York. (Fig. 7.)

### Habitat and ecology

*Capparis xylocarpa* is known from deciduous vine thickets, simple and complex notophyll vine forest and rarely in open woodland.

### Phenology

Flowers have been recorded in August, September and November. Fruit has been recorded in September and December.

### Etymology

The specific epithet *xylocarpa* is derived from the Greek *xylo*-(woody) and - *carpos* (fruit).

### Notes

Seeds are viable and germinate readily when they are firm and brown, within fruits that are hard and green coloured. Whether the fruit ripens from green to orange and dehisces like the similar species *Capparis lasiantha* is unknown. *C. xylocarpa* fruit does not possess suture lines and is therefore unlikely to dehisce.

### Specimens examined

AUSTRALIA. QUEENSLAND. Black Hill Southern Scrub, 15km NE of Kalpowar homestead, 18 Nov. 2014, *S.L.Thompson et al. SLT14940* (BRI); Bathurst Head, Princess Charlotte Bay, Aug./Sep. 1980, *R.Buckley 6677* (BRI); Nesbit River near mouth, Silver Plains, 4 July 1997, *P.I.Forster PIF21376, M.C.Tucker & R.Jensen* (BRI, CNS); Taylors Landing, Claudie River, 4 Oct. 2021, *W.E.Cooper 2789 & T.Hawkes* (CNS); Taylors Landing, Claudie River, 25 Dec. 2019, *W.E.Cooper 2614 & R.Jensen* (CNS); Bramwell gravel pit scrub, 28 June 2021, *W.E.Cooper 2768, R.Jensen & D.G.Fell* (CNS, BRI); Wattle Hills, 16 Oct. 2022, *W.E.Cooper 2894, R.Jensen & F.A.Zich* (CNS); Roadside scrub, Peninsula Development Road, Bramwell, 2 Dec. 2020, *W.E.Cooper 2731, R.Jensen, D.G.Fell & T.Hawkes* (CNS); Blue Tongue Scrub, Steve Irwin Wildlife Reserve, 29 Nov. 2020, *W.E.Cooper 2708, R.Jensen, D.G.Fell & T.Hawkes* (CNS, BRI); Bamaga, Cape York, 1 Sep. 1963, *W.T.Jones 2567* (BRI); Roadside near Somerset camping area, 1 Sep. 2019, *W.E.Cooper 2592 & T.Hawkes* (CNS); Roadside near Somerset, 1 Sep. 2019, *W.E.Cooper 2586 & T.Hawkes* (CNS); Bamaga area, around Somerset Homestead, 1 Sep. 1980, *V.Scarth-Johnson 1048A* (BRI).

***Capparis*** sect. ***Busbeckea*** (Endl.) Hook.f.

*Type species: Capparis nobilis* (Endl.) F.Muell. ex Benth.

***Capparis megacarpa*** W.E.Cooper, *sp. nov*.

*Type*: AUSTRALIA. QUEENSLAND. Cook District: Iron Range Research Station, 9 Mar. 2022, *W.E. Cooper 2841 & J. Pritchard* (holo: CNS 155750 [2 sheet + spirit]), iso: 6 sheets to be distributed to BRI, CANB, K, L, MO, S).

*Capparis* sp. A (in part): Hewson in George, A.S. (1982: 221–222)

*Capparis* sp. (Coen L.S.Smith 11862): Jessup in Henderson (1994: 61), (1997: 43), (2002: 42); Jessup in Bostock & Holland (2007: 42), Guymer in Bostock & Holland (2013), (2016) *Capparis* sp. (Coen): Cooper (2004: 110); Guymer in Bostock & Holland (2010: 37

*Capparis* sp. Coen (L.S.Smith 11862) Qld Herbarium

*Capparis* sp. (Iron Range): Cooper & Cooper (2004: 110)

*Capparis* sp. Iron Range (B.Hyland 16395): Centre for Australian National Biodiversity Research (2010), (Zich 2020)

### Illustrations

Cooper & Cooper (2004: 110), Zich e*t al*. (2020)

### Diagnosis

*Capparis megacarpa* is similar to *Capparis splendidissima* W.E.Cooper, but differs from the latter by having indumentum on new growth golden (*v*. whitish); leaf apex with a mucro 0.75– 2 mm long (*v*. 0.5 mm long); fragrance emitted by unfurling flowers is reminiscent of Ethyl Acetate (*v*. sweetly floral); bud diameter *∼*13 mm (*v*. 17–21 mm); outer sepals free and overlapping in bud (*v*. connate and not overlapping); buds smooth and without a nipple (*v*. ribbed and nippled); stamens *∼*125 (*v. ∼*80); stamen length up to 30 mm (*v*. 60–110 mm); floral gynophore 25–27 mm long (*v*. 65–140 mm long); fruit ellipsoid, 110–145 mm long and 55– 105 x 65–86 mm wide (*v*. globose or elongated globose, up to 94 mm long and diameter 75– 85 mm).

### Description

*Liana* to forest canopy with stem diameter to *∼*80 mm. Stems with grey or brownish bark, fissured vertically becoming tessellated on older wood; branches pendulous. *Indumentum* golden, sometimes drying to whitish. *Stipular spines* persistent, paired, recurved, 2–3 mm long on young stems, becoming acicular on older stems and *∼*10 mm long, on juvenile growth they are acicular and *∼* 6 mm long. *Leaves* discolorous; petioles 4–13 mm long, up to 1.75 mm wide, clothed in very short pilose hairs, glabrescent; lamina oblong or ovate-oblong, 70–160 mm long and 30–73 mm wide, leathery, glabrous; base rounded, obtuse, cuneate or slightly cordate; apex acute and mucronulate, mucro 0.75–2 mm long; entire; venation brochidodromous, primary vein depressed on adaxial surface and raised abaxially; secondary veins 11–16 pairs, flush on adaxial surface and slightly raised on abaxial surface, angle of divergence from primary vein *∼*20°, forming loops 2–3 mm from margin; intralateral veins reticulate. *Inflorescence* an axillary or mostly terminal 4–9-flowered raceme, spines may be present; pedicels 30–56 mm long, puberulent, glabrescent. *Flower* buds globose, smooth and not ribbed, diameter to *∼*13 mm, clothed in minute appressed hairs, apex rounded; fragrant on opening at dusk and smelling strongly of Ethyl Acetate and only slightly florally fragrant on disintegration at dawn. *Calyx* globose; sepals 4 (2 each in 2 whorls), free, overlapping, green; outer pair deeply cymbiform, veins slightly evident but not raised and ribbed, 18–19 mm long, diameter *∼*12 mm, puberulent, green, entire; inner pair obovate, flat, *∼*18 mm long and 12 mm wide, glabrous, green, margin minutely fimbriate. *Petals* 4, obcordate, *∼*32 mm long and 24 mm wide, abaxial surface glabrous, adaxially lanulose, white, base attenuate, apex rounded or emarginate, margin lacerate. *Stamens* numerous *∼*125, filaments 23–30 mm long, white with pale pink bases, anthers *∼*3.5 mm long, cream or grey. *Gynophore* 25–27 mm long, glabrous, pink. *Ovary* mostly zygomorphic-ovoid or rarely ovoid, 4–5.5 mm long and 1.7–2 mm wide, glabrous, ribbed, 5-partite; ovules numerous, >100, diameter *∼* 0.4 mm; stigma sessile, diameter *∼*0.75 mm (Fig. 9). *Fruit*: pedicel 36–55 mm long, swollen at base and apex, indumentum minute and appressed; gynophore 18–22 mm long, swollen at base, glabrous; balausta (hard and indehiscent) is ellipsoid, often with an asymmetrical base and a nippled apex, a few irregular vertical ribs present as well as, puckers or warty protuberances, 110–145 mm long and 55–105 x 65–86 mm wide, green, epicarp 5–10 mm thick; mesocarp fleshy, cream or pale apricot coloured, smells fruity but is bitter to taste; seeds numerous > 100, testa brown, 8–10 x 6–8 x 4–5 mm. Germination epigeal. (Figs. 10 & 11).

**Fig. 8.**
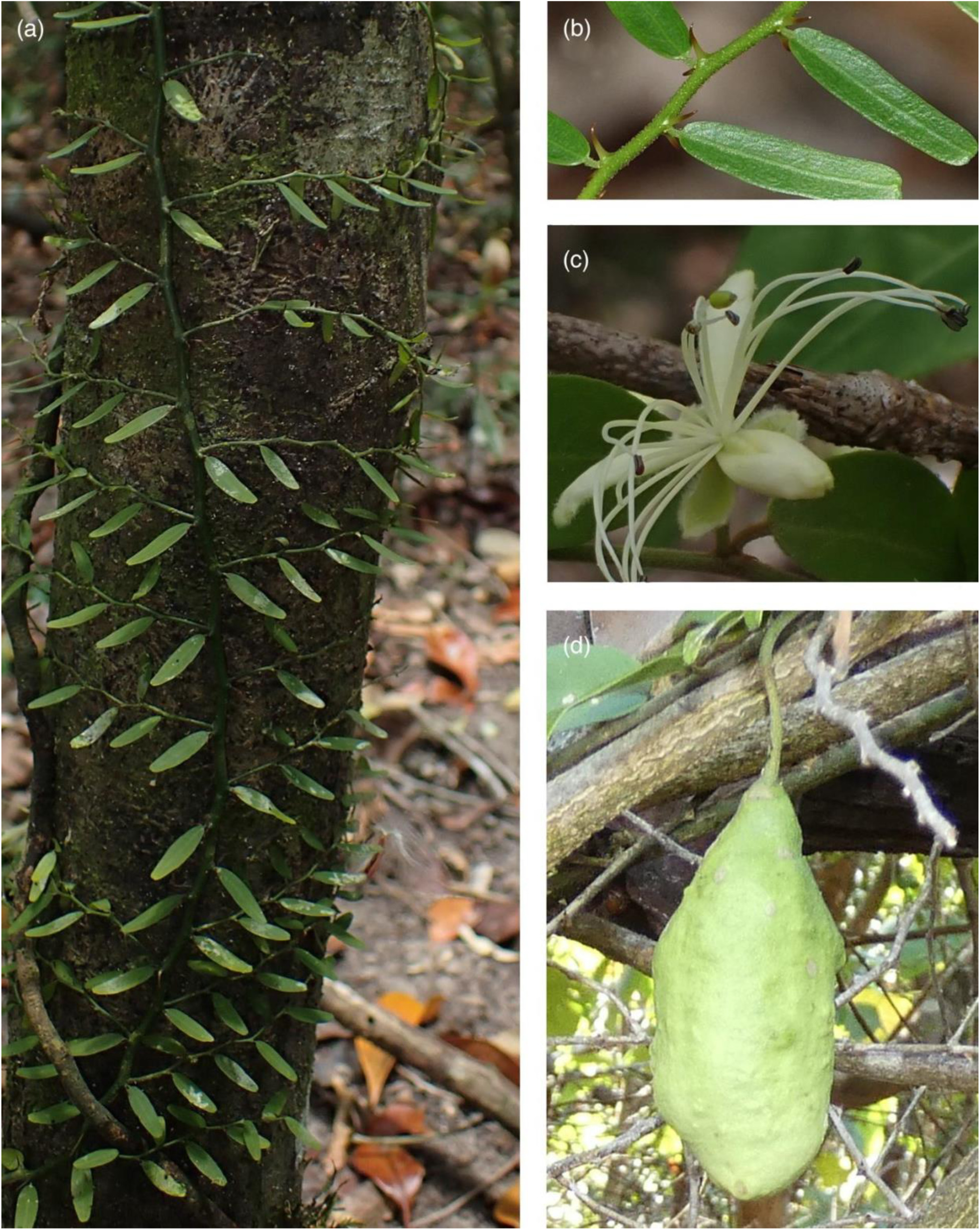
*Capparis xylocarpa*: (*a*) Juvenile growth typically with oblong leaves and stipules hooking onto tree bark. (*b*) Juvenile leaves and stipules. (*c*) Flower at anthesis. (*d*) Fruit. *Cooper 2585*. Photographs: W. Cooper.

**Fig. 9.**
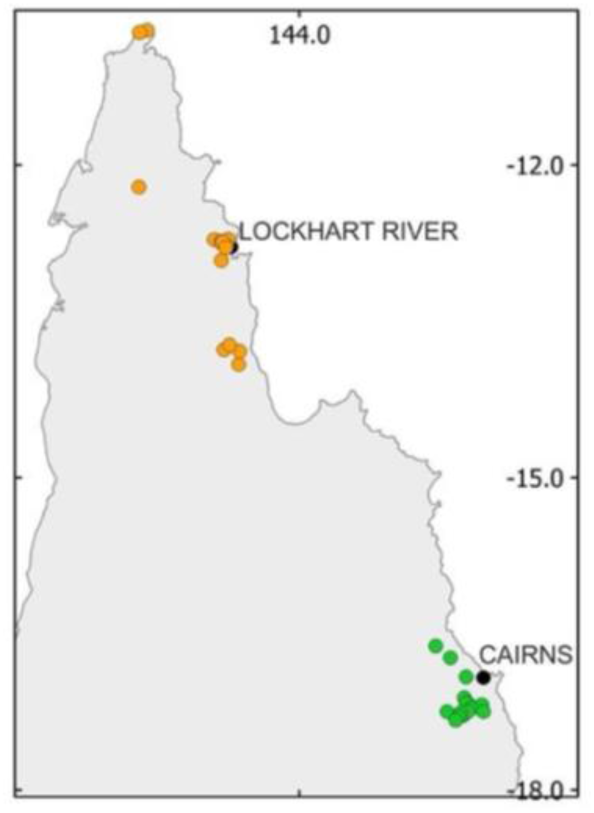
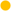 *Capparis megacarpa* and 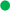 *Capparis splendidissima* distribution.

**Fig. 10.**
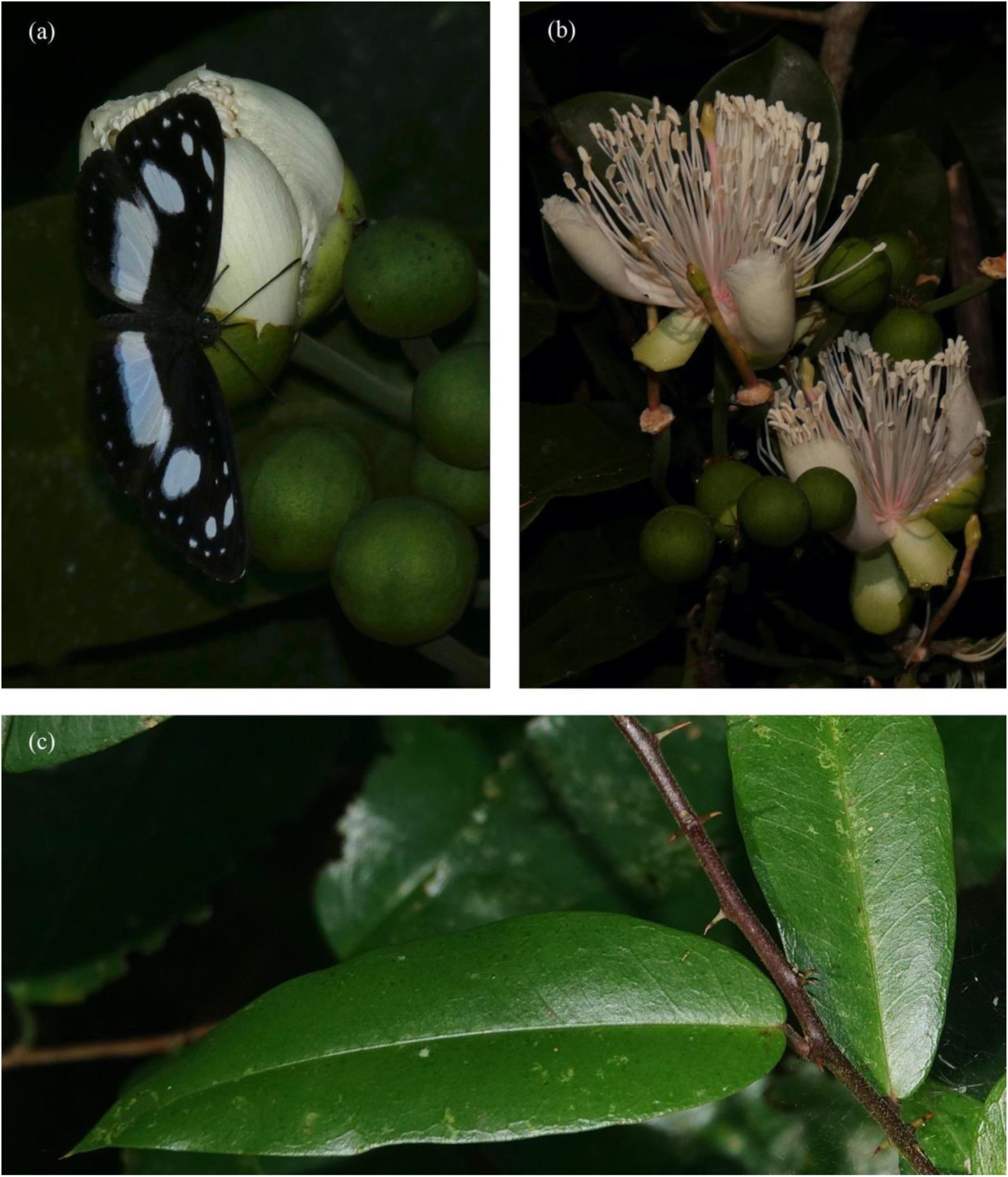
*Capparis megacarpa*: (*a*) Black-eyed Aeroplane (*Pantoporia venilia*) feeding on nectar oozing from an unfurling flower bud. (*b*) Freshly unfurled flowers oozing nectar. (*c*) Juvenile leaves and stipular spines. *Cooper 2841.* Photographs: W. Cooper.

**Fig. 11.**
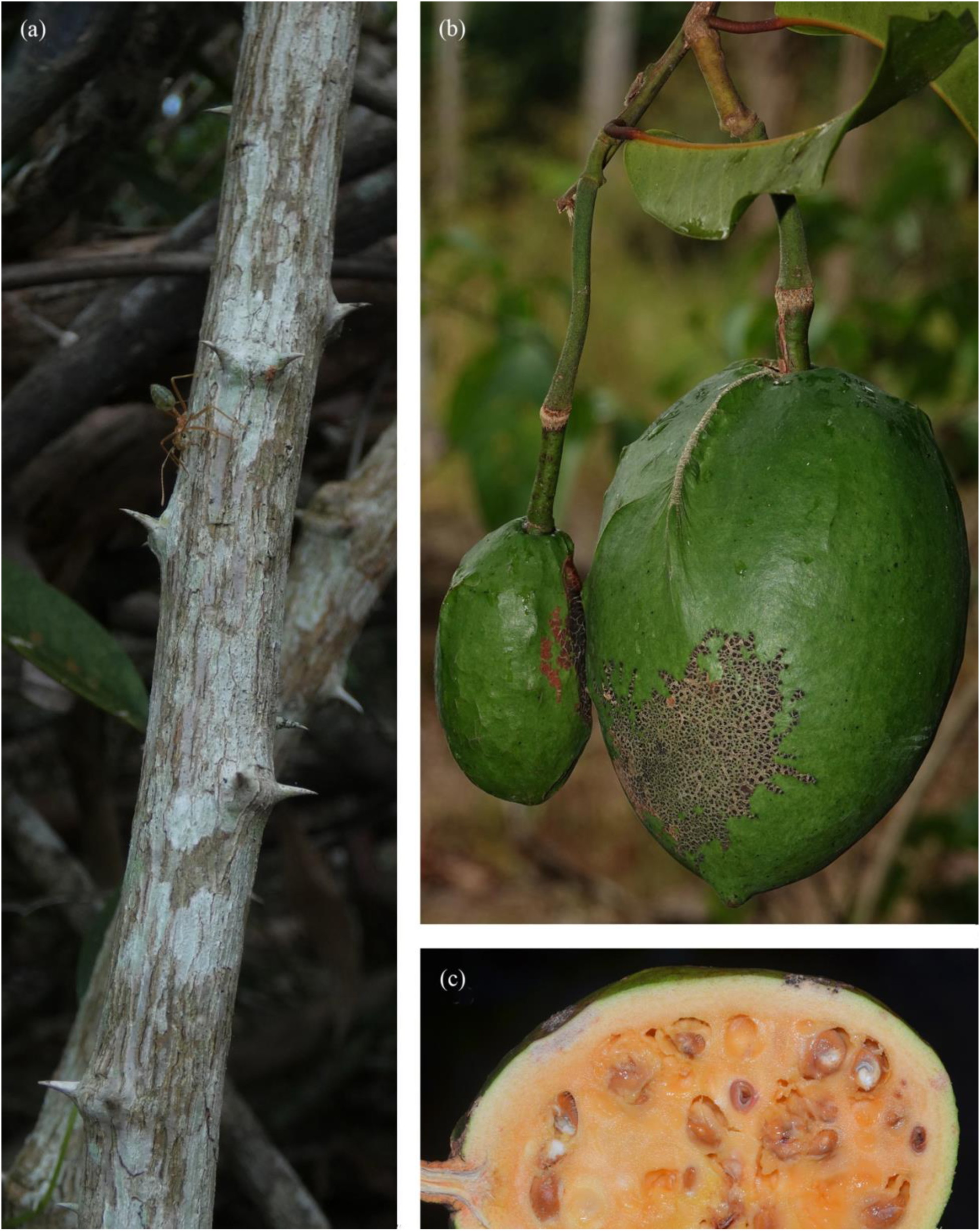
*Capparis megacarpa*: (*a*) Older stems and stipular spines. (*b*) Unripe and ripe fruit. (*c*) Ripe fruit cross-section. *Cooper 2875*. Photographs: R. Jensen.

### Distribution

*Capparis megacarpa* occurs on Cape York Peninsula north from the McIlwraith Range to Cape York, mostly on the eastern side of the peninsula with one record from the central area at Steve Irwin Wildlife Reserve (Fig. 8).

### Habitat and ecology

*Capparis megacarpa* occurs on metamorphic and sedimentary soils as well as sand dunes within various forms of rainforest ranging from well-developed evergreen rainforest, semi-deciduous rainforest, deciduous vine thicket, littoral rainforest, disturbed rainforest regrowth and rarely in woodland.

### Phenology

Flowers have been recorded in March, June and October; ripe fruit has been collected in October and November.

### Etymology

The specific epithet *megacarpa* is derived from the Greek *megas* (great or large) and *karpos* (fruit), referring to the especially large fruit, possibly the largest of any *Capparis* species worldwide.

### Notes

*Capparis megacarpa* is within *C*. sect. *Busbeckea,* which has been suggested to possess connate sepals (Jacobs 1965). However, *C. megacarpa* differs in this morphology by having the outer pair of sepals free and overlapping. This may be relevant to the non-monophyly inferred in this paper.

During the late afternoon prior to flowers unfurling, buds on *Capparis megacarpa* drip copious nectar from between the sepals, attended by numerous insect species.. A different suite of insect species attend the open flowers after nightfall. The flowers are only open for one evening, beginning to disintegrate at dawn.

This species potentially possesses the largest *Capparis* fruit worldwide, weighing up to 395g and with a diameter up to 105 mm.

### Specimens examined

AUSTRALIA. QUEENSLAND. Upper Massey Creek *ca*. 15 miles a little S of ENE of Coen, 13 Oct. 1962, *L.S.Smith 11862* (BRI); North western fall of McIlwraith Range at head of Peach Creek, 1 Oct. 1969, *L.J.Webb & J.G.Tracey 9822* (BRI); Taylors Landing, Claudie River, 4 Oct. 2019, *W.E.Cooper 2790 & T.Hawkes* (CNS); Max’s Ridge, forest edge beside old quarry, 27 Feb. 2022, *W.E.Cooper 2838 & J.Pritchard* (CNS); Max’s Ridge, forest edge beside old quarry, 11 Oct. 2022, *W.E.Cooper 2875, J.Pritchard*, R.*Jensen & F.A.Zich* (CNS); Max’s Ridge, forest edge beside old quarry, Nov. 2022, *W.E.Cooper 2937 & J.Pritchard* (CNS); Claudie River, 2 June 2000, *B.Hyland 16395* (CNS); West Claudie roadside, 26 Dec. 2019, *W.E.Cooper 2623 & R.Jensen* (CNS); West of Tozer Gap, 29 Oct. 1999, *B.Hyland 21382V* (CNS); Young Ones Track, roadside, Iron Range National Park, 12 Oct. 2016, *W.E.Cooper 2362* (CNS); SIWR, Moaning Scrub, 4 June 2022, *W.E.Cooper 2855 & T.Hawkes* (CNS); Chili Creek, 28 Oct. 1999, *B.Hyland 16300* (CNS); Wattle Hills, 1 Sep. 2019, *W.E.Cooper 2799 & T.Hawkes* (CNS); Wattle Hills, 4 Apr. 2022, *W.E.Cooper 2848A & H.Mara* (CNS); Table Range, Dead Horse Creek, 23 Oct. 1973, *A.W.Dockrill 765* (CNS); 1.8 km SE of Punsand Bay camp, Cape York, 25 July 2018, *D.G.Fell DGFIRRS245* (BRI, CNS); Evans Bay, Cape York Peninsula, 29 July 2018, *D.G.Fell DGFIRRS236* (BRI, CNS).

***Capparis loxophleba*** W.E.Cooper & Joyce, *sp. nov*.

*Type*: AUSTRALIA. QUEENSLAND. Wide Bay District: Mary Cairncross Scenic Reserve, Blackall Range, 3km SE of Maleny, 10 Jan. 2008, *P.I. Forster PIF33222 G. Guymer & P. McAdam* (holo: BRI 43409; iso: CNS, MEL, NSW).

### Illustrations

Leiper *et al*. (2008, p.355 as *Capparis arborea* [hairy-leaved form])

### Diagnosis

*Capparis loxophleba* is similar to *C. arborea* (F.Muell.) Maiden but differs from the latter by the habit of being a liana or scrambler (*v*. shrub or tree); stipular thorns to 2 mm long (v. ∼7 mm long); secondary veins 10–16 pairs (v. 8–12); secondary vein angle of divergence from primary vein 70–85° (*v*. 40–60°); leaf hairs persistent on abaxial surface especially along the midrib (*v*. glabrous).

### Illustrations

Leiper, Glazebrook, Cox & Rathie (as *Capparis arborea* hairy-leaved form) (2008:355)

*Shrub or liana* to canopy. Bark rough with swollen horizontal bands on larger stems.

*Stipular spines* acicular, up to 2 mm long, mostly absent from new stems. *Indumentum* slightly tortuous, cream or yellowish. *Leaves* discolorous; petioles 4–10 mm long, 1–1.6 mm wide, clothed in slightly tortuous cream or yellowish hairs; lamina oblong, 40–125 mm long and 20– 53 mm wide, leathery, adaxially glabrous with the exception of short tortuous cream or yellowish hairs along the midrib, abaxially with short tortuous cream or yellowish hairs especially near base and along primary vein; base cuneate or rarely obtuse or rounded; apex acute and mucronulate, mucro up to 1.5 mm long; entire; venation brochidodromous, primary vein depressed on adaxial surface and raised abaxially; secondary veins 10–16 pairs, flush on adaxial surface and slightly raised on abaxial surface, angle of divergence from primary vein 70–85°, forming loops 3–5 mm from margin; intralateral veins reticulate. *Inflorescence* an axillary pair or solitary flower or a terminal raceme; pedicels 35–60 mm long, clothed in short tortuous cream or yellowish hairs; buds ovoid, smooth and not ribbed, sparsely clothed in tortuous cream or rusty hairs, apex beaked, diameter ∼ 12 mm. *Flowers* fragrant; sepals 4 (2 each in 2 whorls), connate, green; outer pair deeply cymbiform, apex thickened and squarish, smooth and not ribbed, *∼* 15 mm long, diameter *∼* 15 mm, puberulent, green, entire; inner pair cymbiform, obovate, 12–14 mm long and 8–10 mm wide, abaxially and adaxially sericeous, green, margin minutely fimbriate. *Petals* 4, obcordate, *∼* 30 mm long and 18 mm wide, both sides lanulose especially towards basal half, white, base attenuate, apex rounded or emarginate, margin lacerate. *Stamens* numerous, filaments *∼*35 mm long, white, anthers *∼*2 mm long, grey; gynophore *∼*30 mm long, thinly lanulose in lower section. *Ovary* spindle-shaped, *∼* 4.5 mm long and *∼* 2.25 mm wide, glabrous, ovules numerous; stigma sessile, diameter *∼* 0.5 mm. *Fruit*: pedicel *∼* 30 mm long, gynophore 30–40 mm long, swollen at base, thinly lanulose near base; balausta (hard and indehiscent) globular often with a nippled apex, diameter 28–40 mm, smooth, green; seeds numerous (Fig. 12).

### Distribution

*Capparis loxophleba* occurs in SE Qld from the Border Ranges to Wrattens National Park west of Gympie south to the Macleay River NSW (Fig. 11).

**Fig. 12.**
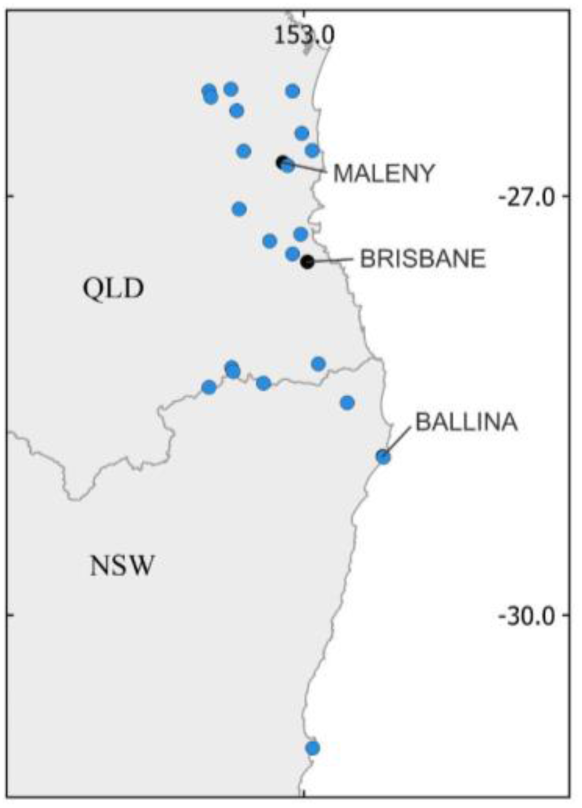
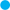 *Capparis loxophleba* distribution.

**Fig. 13.**
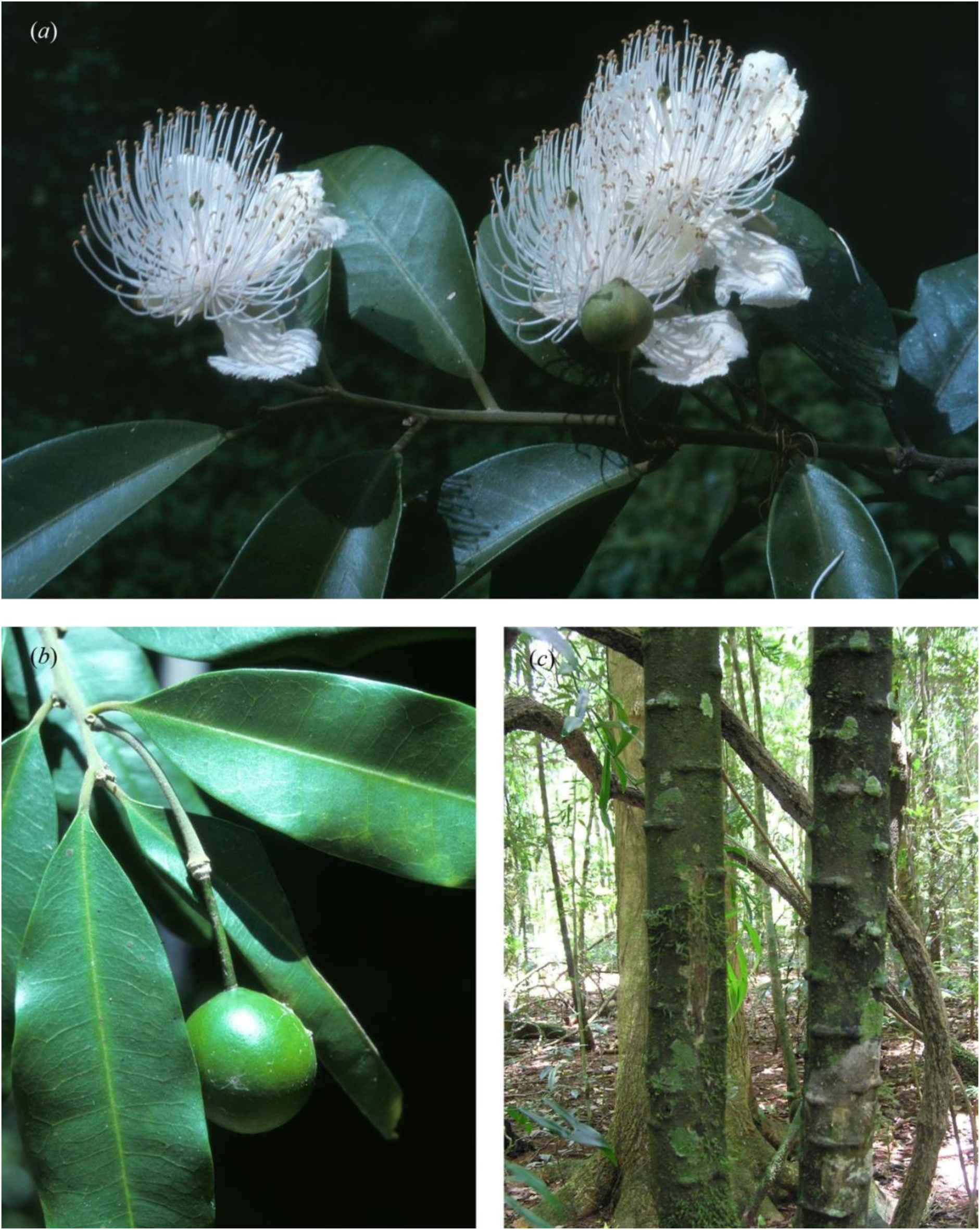
*Capparis loxophleba*: (*a*) Bud, flowers and leaves. (*b*) Leaves with widely angled lateral veins and fruit. (*c*) Large stems with rough bark and swollen horizontal bands. Photographs: (*a*) & (*b*) G. Leiper, (*c*) W. Cooper.

### Habitat and ecology

*Capparis loxophleba* occurs in complex notophyll vine forest.

### Phenology

Flowers have been recorded in January and February and fruit has been recorded for September.

### Etymology

The specific epithet *loxophleba* is derived from the Greek *loxos* (slanting, oblique or cross wise) and *phlebius* (-veined) referring to the lateral veins occurring at a wide angle to the primary vein.

### Specimens examined

AUSTRALIA. QUEENSLAND. State Forest 639 Wrattens, Beauty Spot 50, end of Snake Road, 1 Mar. 2003, *P.I.Forster PIF29244 & M.C.Tucker* (BRI); Fireclay Scrub, Wrattens State Forest, 4 Feb 1988, *W.J.McDonald 4162* (BRI); Mary Cairncross Scenic Reserve, Blackall Range, 4 Dec. 2004, *P.I.Forster PIF30428* (BRI, MEL); NW slopes of Mt Glastonbury, 30 Dec. 1991, *P.I.Forster PIF9274 & P.R.Sharpe* (BRI); Mt Lindsay, 16 Oct. 1992, *P.I.Forster PIF12183 & G.Leiper* (BRI); North Pine River, Petrie, 18 miles north of Brisbane, 4 Jan. 1932, *S.T.Blake 3141* (BRI); Kin Kin, Wide Bay, Dec. 1919, *W.D.Francis s.n.* 28029 (BRI); Collins Road, Yandina, slopes of Mt Ninderry, Dec. 1993, *Thomas 53* (BRI); Mount Glorious Road, driveway into ‘Skye’ property, *∼* 0.25km S of The Summit, Mount Glorious, 25 Nov. 2007, *S.P.Phillips 1845 & B.A.Phillips* (BRI); Nineteen Logging area, Timber Reserve 209, Mount Brisbane, 21 June 1990, *P.I.Forster PIF6868* (BRI, CNS); Sunday Creek Road, Conondales, 9 Jan. 2002, *P.I.Forster PIF28100 & G.Leiper* (BRI, MEL, NSW); Portion 17V Cainable, Lamington National Park, 29 June 1978, *W.J.McDonald 2126, L.W.Jessup & W.G.Whiteman* (BRI); SW slope of Wilsons Peak, 27 Jan. 1990, *P.I.Forster PIF6220 L.H.Bird & P.D.Bostock* (BRI); The Head track to Lizard Point, Main Range National Park, 17 Dec. 2004, *P.I.Forster PIF30494, R.Jensen & M.C.Tucker* (BRI); Gold Creek, Brookfield, 1977, *J.N.Hargreaves JT696* (BRI); South Amamoor Creek headwaters, Amanmoor Range, 20 May 1994, *P.I.Forster PIF15211* (BRI); Martins Creek, Lindsay Road Buderim, Jan. 2004, *G.Leiper s.n.* (BRI); North Pine River, Petrie, 18 miles N of Brisbane, 4 Jan. 1932, *S.T.Blake 3141* (BRI); One Tree Hill, Mount Coot-tha, Brisbane, Simmonds, Aug. 1960, *Simmonds s.n.* (BRI) 2217; Tambourine, July 1915, *J.F.Shirley s.n.* (NSW). NEW SOUTH WALES: Acacia Creek [(NSW)] near Killarney ([Qld]), Sep. 1905, *W.Dunn s.n. NSW 405645* (BRI, NSW);.Tintenbar, Dec. 1895, *W.Baeuerlen s.n.* (NSW); Ballina, Dec. 1891, *W.Baeuerlen 659* (NSW); Macleay River, Sep. 1987, *leg. ign.* (NSW 405646); Gloucester, Bundook Road, *c*. 2.5 km directly WSW of Bundook, 16 Dec. 2013, *R.W.Purdie 9290,* (CANB, NSW); 5.1 km on Cudgera Creek Road from Old Pacific Highway at Mooball, 11 Feb. 2014, *R.L.Johnstone 3450, G.Errington & K.R.Kupsch* (CANB, NSW).

*Capparis splendidissima* W.E.Cooper, *sp. nov*.

*Type:* AUSTRALIA. QUEENSLAND. Cook District: Carpark at Curtain Fig National Park, 5 Jan. 2021, *R.Jensen 4341 & W.Cooper* (holo: CNS 152698 [2 sheets + 1 spirit], iso: 7 sheets to be distributed to BRI, CANB, DNA, K, L, MEL, MO).

*[Capparis ornans auct. non* F.Muell. ex Benth.: sensu Hyland *et al*. in Cooper & Cooper (1994: 302), Cooper & Cooper (2004: 110), Zich *et al*. 2020]

### Illustrations

As *Capparis ornans*: Cooper & Cooper (2004: 110), Zich *et al*. 2020.

### Diagnosis

*Capparis splendidissima* is similar to *C. ornans* F.Muell. ex Benth. but differs from the latter by the lack of indumentum on new leaf laminas (*v*. densely hairy); senescing leaves green (*v*. red or reddish); leaf shape oblong (*v*. ovate, broadly elliptic or subrhombic); leaf width equal to less than half (usually about 1/3) of the length (*v*. equal to half or greater than half the length); leaf apex with a mucro (*v*. mucro absent); leaf lateral veins 8–13 pairs (*v*. 6–7 pairs); flower pedicel 80–120 mm long (*v*. up to 42 mm long); floral gynophore 90–120 mm long (*v*. 65–76 mm long); flower buds ovoid and distinctly ribbed (*v*. globose and not distinctly ribbed); stamens 80–100 mm long (*v*. less than 75mm long); fruit gynophore 90–120 mm long (*v*. 60–75 mm long); ovary 5–6 mm long (*v*. 10 mm long). Fruit shape orbicular and not ribbed (*v*. ovoid ellipsoid, often slightly ribbed towards base); distribution on the Atherton Tablelands and general area from Millaa Millaa to Julatten (*v*. south from the Rockingham Bay/Tully area to central Queensland).

### Description

*Liana* to forest canopy (or shrubby when small or a plant isolated by forest clearing), stem diameter to 80 cm, branches pendulous; bark brown, rough and fissured vertically. *Indumentum* fawn or whitish. Young branches clothed in simple prostrate hairs. *Stipular spines* paired, mostly present on non-flowering stems, recurved, decurrent, 2.5–6.5 mm long, spines on juvenile growth acicular and up to 10 mm long. *Leaves* discolorous; petioles 3–30 mm long, 2.25–2.5 mm wide, puberulent; lamina discolorous; oblong, ovate-oblong or rarely elliptic, 50– 160 mm long and 20–72 mm wide, coriaceous, glabrous, base cuneate or rounded rarely slightly cordate, apex acute or obtuse, often mucronulate, mucro *∼*0.5mm long, entire, oil dots numerous, round or slightly elongated; venation brochidodromous, primary vein ± flush or shallowly depressed on adaxial surface and raised abaxially; secondary veins 8–13 pairs, slightly raised on both sides, angle of divergence from primary vein 20–35°, forming loops 2– 4 mm from margin; intralateral veins reticulate. *Inflorescence* a terminal 1–5-flowered supra-axillary row or a solitary flower in upper axils, spines rarely present on flowering stems; pedicels 50–120 mm long, puberulous, terete when fresh and ribbed when dried;. *Flower* buds ovoid with a nippled apex, diameter 17–21 mm, ribbed, clothed in appressed hairs; flowers pleasantly fragrant. *Calyx* actinomorphic; sepals 4 in two whorls, outer pair initially connate, separating into 2 cymbiform sepals, ribbed, 17–25 mm long and 15–18 mm wide, abaxially puberulous becoming glabrescent, adaxially glabrous, dark green; inner pair free, oblong, shallowly cymbiform, smooth, with a central rib, 18–23 mm long and 10–11 mm wide, glabrous abaxially, lanulose on adaxial surface, pale green, margin minutely fimbriate and often with an abruptly acute apex. *Petals* 4, flabellate, 25–55 mm long and 13–28 mm wide, abaxial surface glabrous distally and sparsely lanulose towards base, adaxially lanulose, white, apex fimbriate and deeply incised. *Stamens* numerous *∼*80, filaments 60–110 mm long, sparse hairs towards base, white, anthers 3–4 mm long, cream-coloured. *Gynophore* 65–140 mm long, mostly glabrous with a few sparse erect hairs towards base, palest pink. *Ovary* narrowly ovoid, shallowly ribbed, 4-partite, 5–6 mm long and 2.5 mm wide, glabrous, green; ovules several; stigma sessile, globular, diameter *∼*1 mm (Fig. 13). *Fruit*: pedicel up to 90 mm long, glabrous or a few minute hairs may be present; gynophore up to 150 mm long, glabrous; balausta (hard and indehiscent) often with a sparsely puckered or warty epicarp, globose or elongated-globose, up to 94 mm long, diameter 75–85 mm, glabrous, dark green to blackish; pulp surrounding seeds copious, creamy-yellow, sickly sweet smelling; seeds *∼*75, brown, asymmetrically reniform, 12–15 x 9–11 x 5–6 mm (Fig. 13). Germination epigeal. *Seedling* stems and petioles clothed in crisped hairs; stipular spines acicular, up to 10 mm long; lamina ovate, 35–57 mm long and 14–30 mm wide, adaxially glabrous, abaxially with crisped hairs along midrib; base cordate or rounded, apex acute with a mucro 1.5–2.5 mm long. (Fig. 13)

**Fig. 14.**
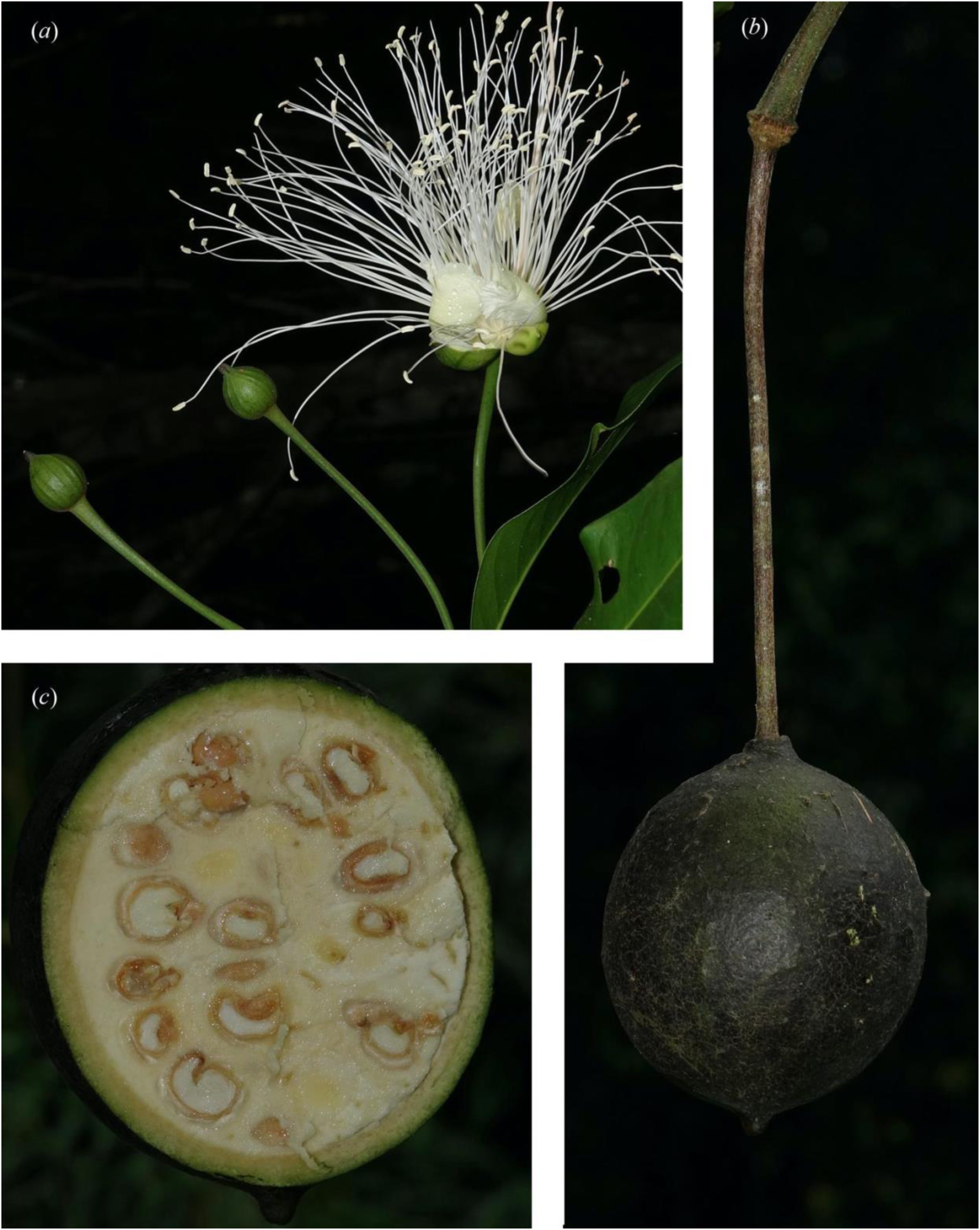
*Capparis splendidissima*: (*a*) Buds and flower at anthesis, *Cooper 2621*. (*b*) Ripe fruit hanging on gynophore. (*c*) Fruit cross section. *Jensen 4341*. Photographs: R. Jensen.

### Distribution

*Capparis splendidissima* is recorded on basalt and granite soils in well-developed rainforest in the Wet Tropics bioregion in north Qld from the Millaa Millaa area to Mossman Gorge, at altitudes from 80–1000 m (Fig. 8).

### Habitat and ecology

*C. splendidissima* grows in semi-deciduous and evergreen rainforest.

### Phenology

Flowers have been recorded in December and January and fruit in January and February.

### Etymology

The specific epithet *splendidissima* is from the Latin *splendidus* (brilliant, splendid, showy) and -*issimus* (superlative), referring to the bounty of spectacular large fragrant white flowers.

### Notes

*Capparis splendidissima* has long been included within the concepts of *Capparis ornans* F.Muell. ex Benth. We present evidence that this tropical rainforest liana possesses significantly different features including the leaves, inflorescences, flowers, fruit, habitat and has a distribution disjunct from *C. ornans,* which occurs in *Eucalyptus* and Brigalow forests to the south of *C. splendidissima*.

*Capparis splendidissima* occurs in rainforest within the Wet Tropics bioregion from Millaa Milla and the Goldsborough Valley north to Mossman Gorge. By contrast,*C. ornans* occurs in Eucalypt woodland and Brigalow communities from the Gladstone area north to the Rockingham Bay.

### Specimens examined

AUSTRALIA. QUEENSLAND: State Forest Reserve 191, 6 Nov. 1978, *B.Gray 20057V* (CNS); State Forest Reserve 310, Goldsborough, 1981, *S.J.Dansie AFO 5221* (CNS); Wooroonooran National Park, near Stallion Pocket off Goldsborough Road, 22 Dec. 2009, *A.Ford 5651b* (CNS, BRI); Lake Barrine, 15 Jan. 1992, *W.Cooper & W.Cooper 131* (CNS); Eastern Connection Road off Powley Road, 12 Dec. 2001, *W.Cooper & W.Cooper 1639* (CNS); Tinaroo, 7 Dec. 1966, *W.E.Volck AFO 4200* (CNS); Fong On Bay area, Danbulla, 20 Feb. 1996, *R.Jensen 596* (CNS); Moseley Road, Barron River, East Barron, 22 Dec. 1994, *G.Sankowsky 1431* (CNS); Curtain Fig National Park, 29 Dec. 2019, *W.E.Cooper 2621 & R.Jensen* (CNS); Curtain Fig carpark, SW of Yungaburra, Jan. 2015, *R.Jensen 3557* (BRI); Allumbah Pocket, Petersen Creek, Yungaburra, 28 Dec. 2001, *R.Jensen 1174* (BRI, L); Pinnacle Pocket road, off Atherton-Yungaburra road, 30 Dec. 2019, *R.Jensen 4215* (BRI); Nature Reserve, 4 km WNW of Yungaburra, 28 Mar. 2017, *M.F.Braby 207* (CANB); State Forest Reserve 185, Downfall Logging Area, SFR 185, 25 June 1981, *B.Gray 20166V* (CNS); State Forest Reserve 607, Shoteel Logging Area, 7 Jan. 1982, *B.Gray 2369* (BRI, CNS); State Forest Reserve 1229, Danbullan, 12 Aug. 1992, *B.Hyland 21250V* (CNS); Ca. 2 km NNW of East Barron, Barron River, 20 Dec. 1994, R.*Jensen 86* (BRI, CNS); Near Flin Creek crossing on Pinnacle Road, *ca.* 10 km ENE of Julatten, 22 Dec. 1976, *V.K.Moriarty 2185* (CNS, BRI); Brooklyn, near water intake at Hunter Creek, Australian Wildlife Conservancy, 23 Jan. 2021, *W.E.Cooper 2746 & R.Jensen* (CNS); Spring Creek Road, 26 Nov. 2003, *R.L.Jago 6600* (BRI); Mossman Gorge National Park, Jan 1992, *R.Russell s.n.* (BRI).

***Capparis bancroftii*** (C.T.White ex. M.Jacobs) W.E.Cooper & Joyce, *comb. et stat. nov. Capparis loranthifolia* var. *bancroftii* C.T.White ex M.Jacobs, *Blumea* 12(3): 519 (1965).

*Type citation*: *”T.L. Bancroft s.n.* (BRI, sh. 33,475) Australia, Queensland, Eidsvold, fr.”. *Type*: AUSTRALIA. QUEENSLAND. Burnett District: Eidsvold, *s.dat.*, *T.L. Bancroft s.n.* (*holo*: BRI 33259 [image!]).

### Illustrations

As *Capparis loranthifolia*: Harden *et al* (2025: 9, 15, 918SP).

*Capparis bancroftii* differs from *Capparis loranthiifolia* by leaves coriaceous (v, chartaceous); leaf bases cuneate or rounded (*v*. long attenuate); secondary veins 5–10 distinct pairs (*v*. 3–6 indistinct pairs); reticulate venation distinct (*v*. indistinct) (Fig. 14). The protuberances on the fruit surface of *C. bancroftii* have previously been described as a diagnostic feature for this species. However, this texture, which is not consistent on all specimens within this species may be caused by insect parasitism (potentially wasps) (Fig. 14). Warty, echinate or muricate surfaces are often present in the fruit of many *Capparis* species. DNA analysis has shown *C. bancroftii* to be more closely related to *C. arborea* than to *C. loranthifolia* Lindl.

**Fig. 15.**
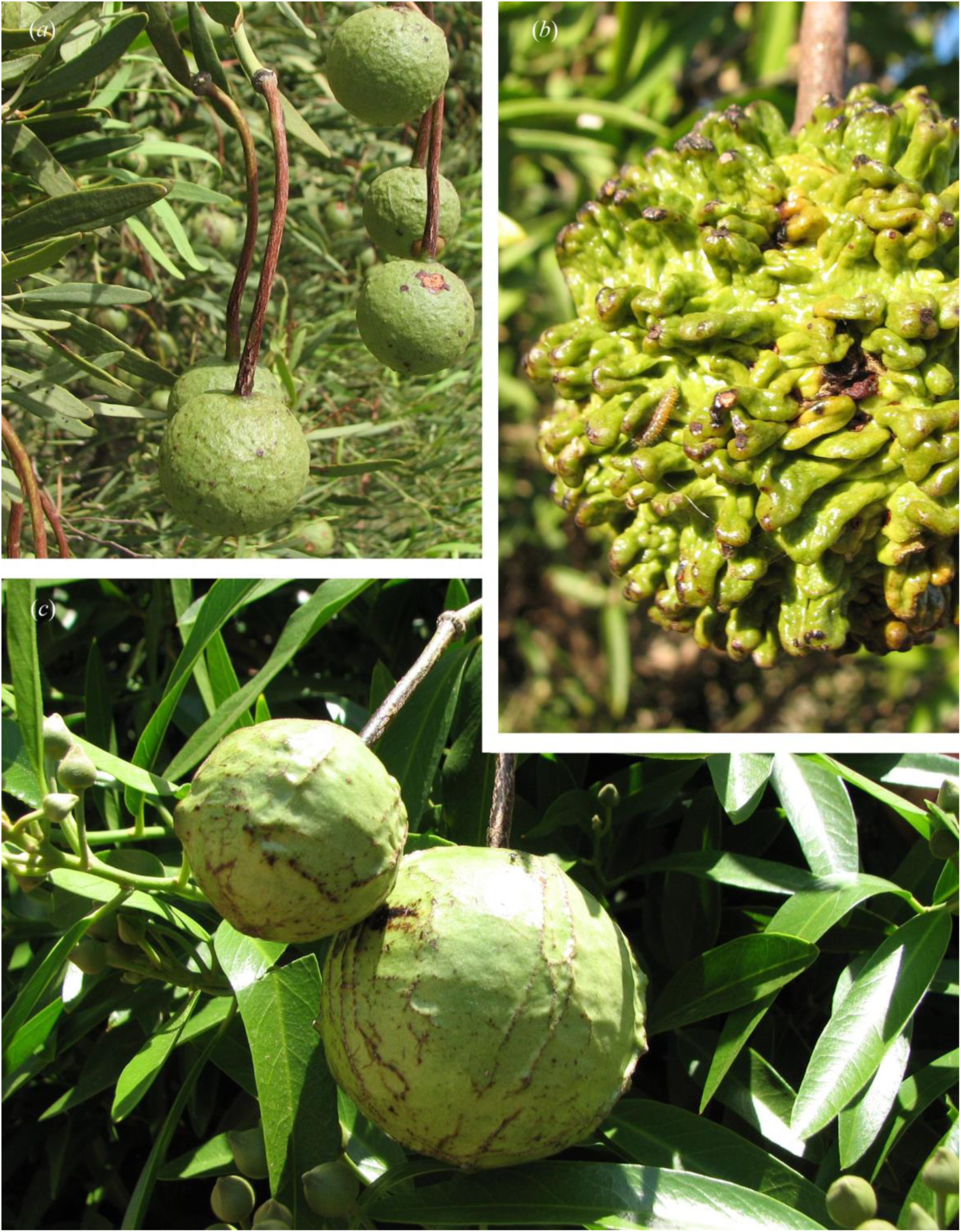
*Capparis loranthifolia*: (*a*) Leaves and fruit, unvouchered. *Capparis bancroftii*: (*b*) Fruit with densely warty surface and freshly emerged caterpillar (potentially the cause of the protuberances), unvouchered. *(c)* Leaves and fruit, fruit surface with scattered small protuberances, typical of many *Capparis* fruit, unvouchered. Photographs: C. Eddie.

### Etymology

The specific epithet *bancroftii* is in honour of Australian physician and naturalist Thomas Lane Bancroft (1860–1933).

### Key to *Capparis* in Australia using mature growth

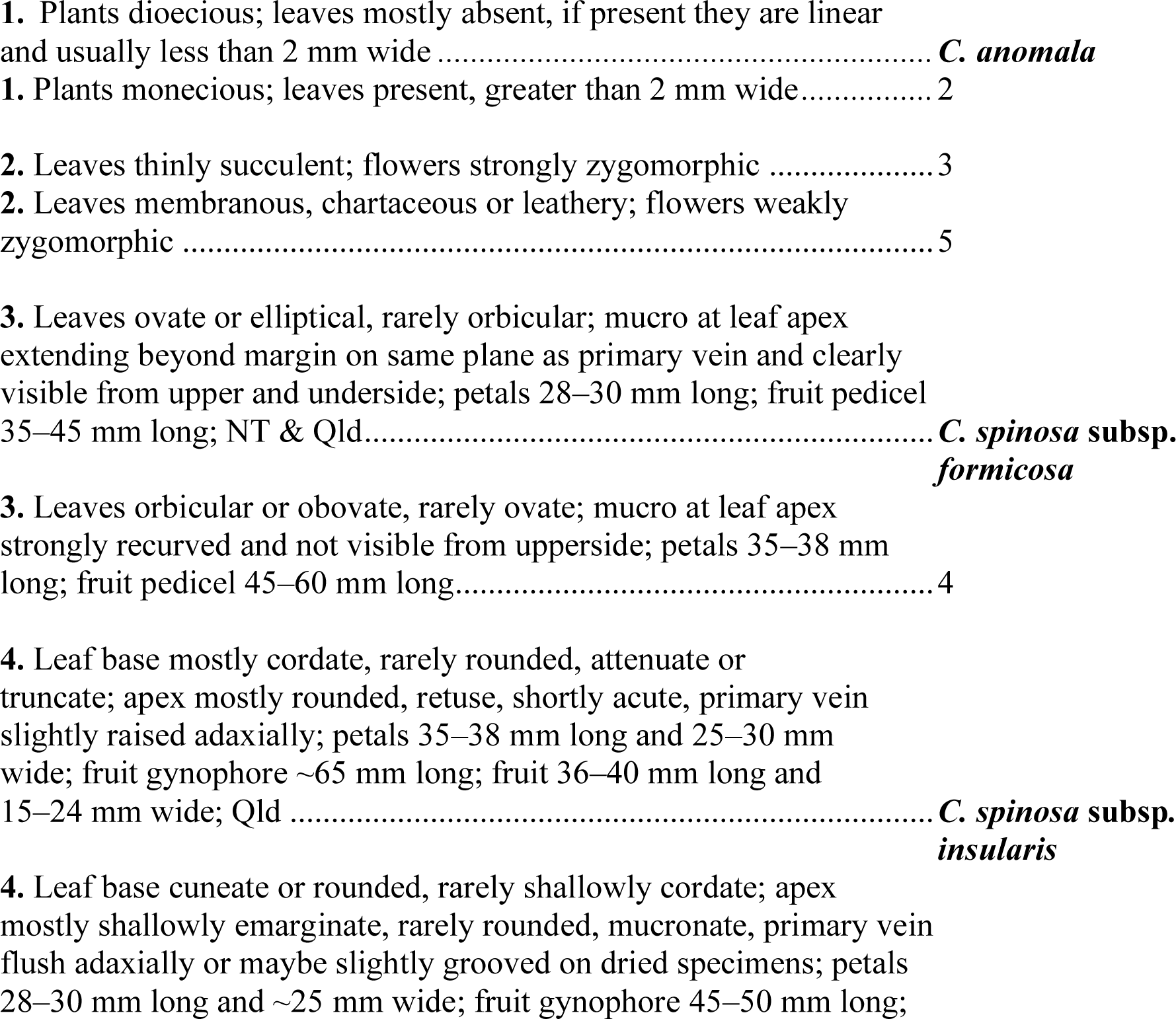
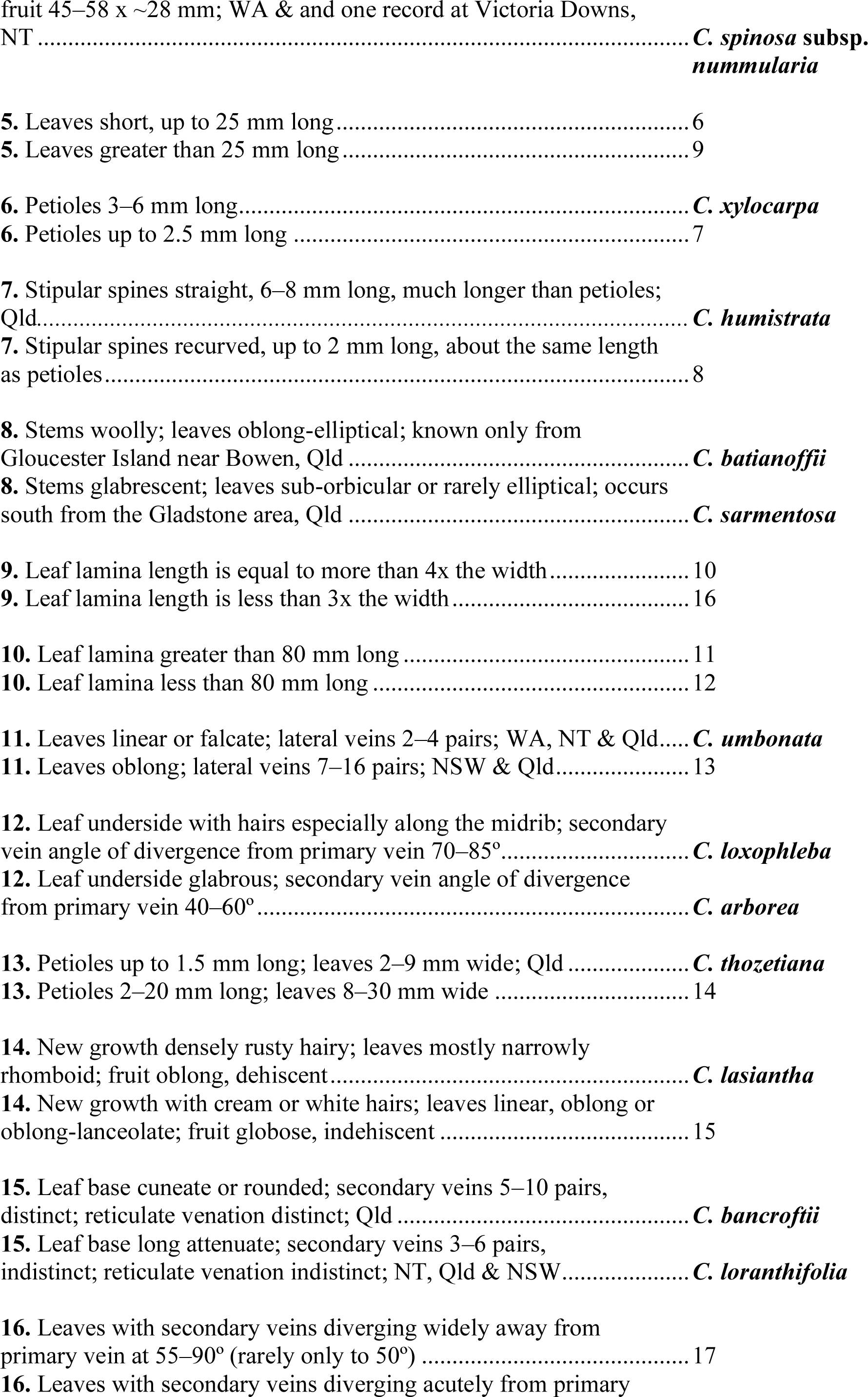
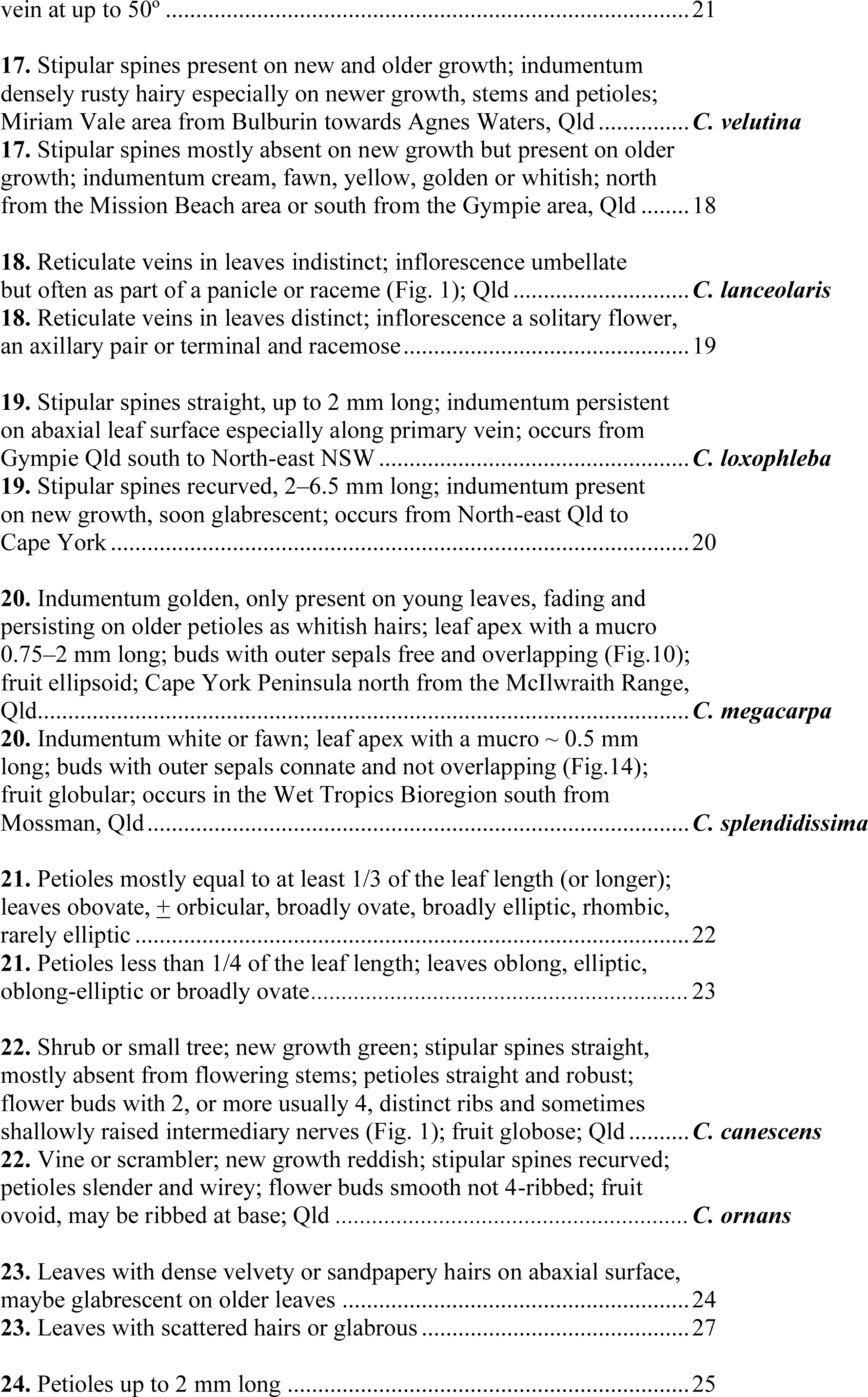
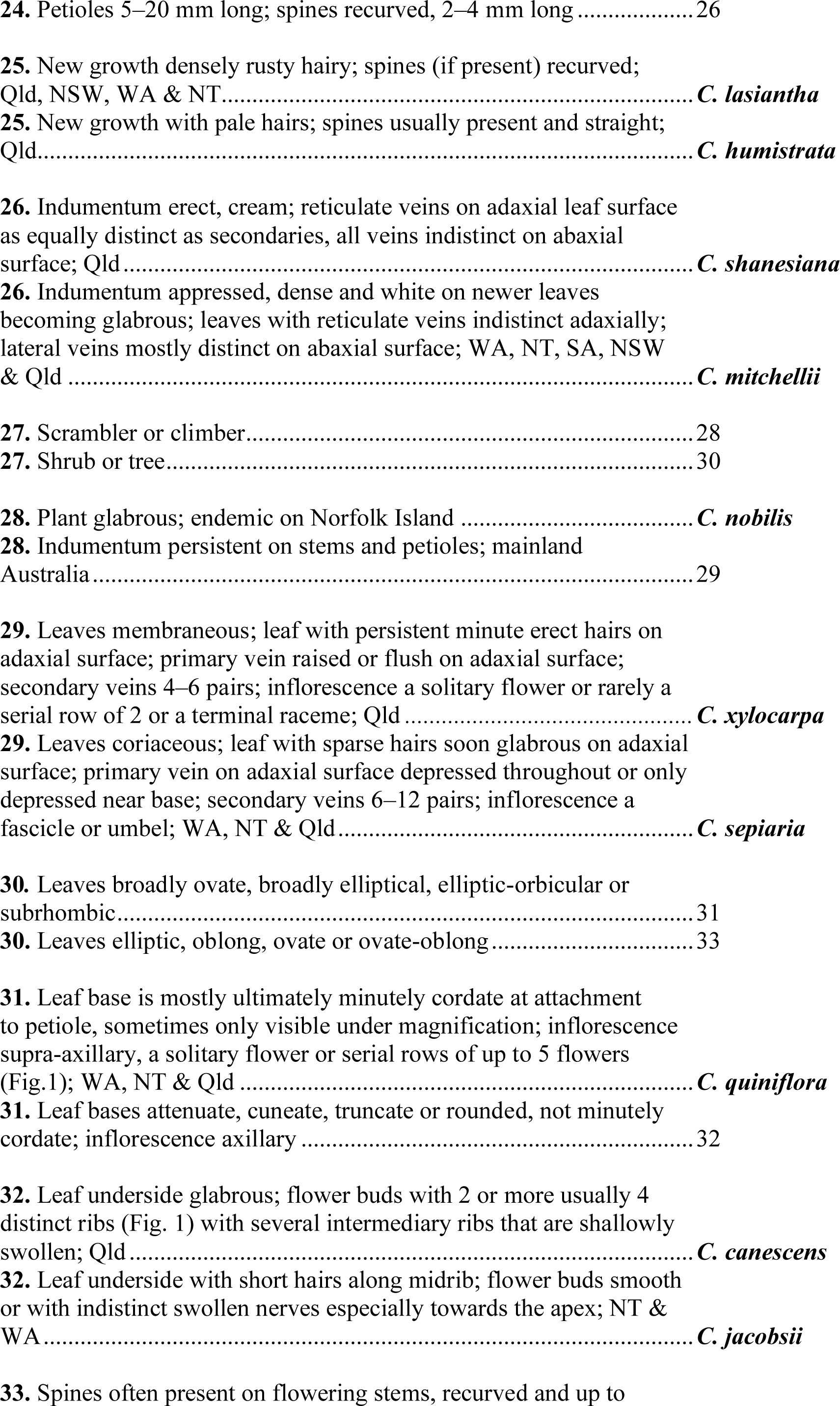
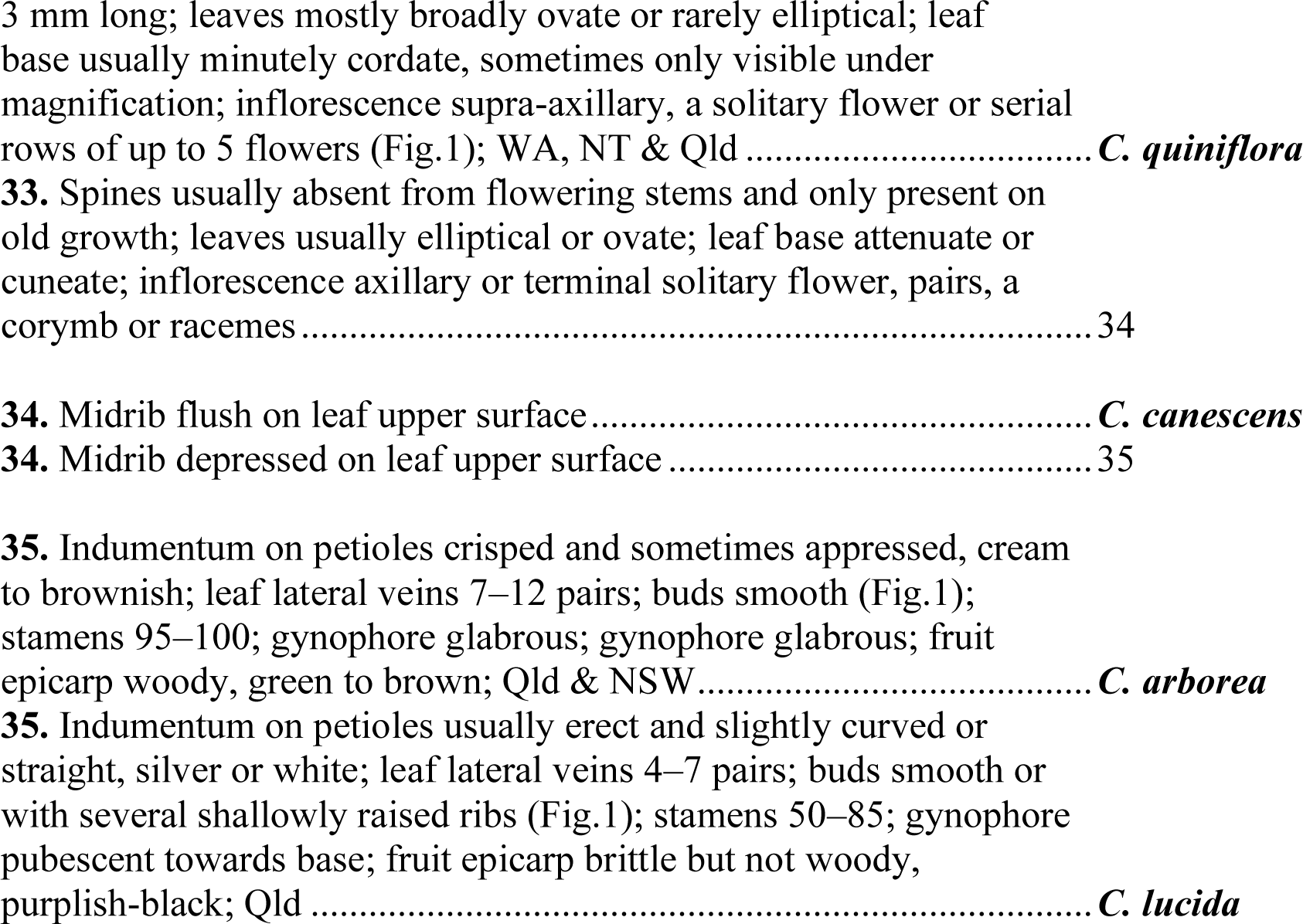

## Acknowledgements

Thanks to Tim Hawkes for much assistance with collecting flowering and fruiting specimens of *Capparis xylocarpa,* Rigel Jensen for *C. splendidissima,* John Pritchard and Bianca op den Brouw for *C. megacarpa* and Bryony Barnett for *C. spinosa* subsp. *formicosa.* For *C. loxophleba* Paul Forster, Glenn Leiper, Spencer Shaw and Stephanie Reif were all most helpful in finding plants in the field. Craig Eddie, Rigel Jensen, Glenn Leiper and Tony Start are thanked for their excellent photos. Thanks also to Anita and Keith Cook at Iron Range Research Station (IRRS) for their generous hospitality and permission to collect *C. megacarpa* on their property. Angharad Johnson and Eugenia Pacitti MEL), Nigel Fechner, Paul Forster and Andrew Franks (BRI), Nick Cuff and David Albrecht (DNA) are thanked for scans and other herbarium assistance. PERTH, BRI, CANB & MEL are thanked for voucher loans. Seedlings were germinated by John Pritchard (*C. megacarpa*), Kylie Freebody (*C. splendidissima*) and Gaylene Sheather (*C. xylocarpa*) and Steve Murphy created magnificent maps. Thank you also to Alina Höwener for assistance in the laboratory, to Gordon Guymer for useful comments on an earlier version of the manuscript, and to two anonymous reviewers whose feedback greatly improved the manuscript. Plant material was collected under permits issued to Darren Crayn by the Queensland Government.

## Data availability statement

Data used in this study can be accessed via NCBI’s GenBank; accession numbers for all data can be found in Supplementary Table 1. Treefiles can be found at Zenodo.

