## Supplementary Material for "The complex evolutionary history of *Capparis* L. (Capparaceae Juss.) in Australia, and the description of five new species and two new subspecies"

1. List of samples
2. Summary of data for each sample
3. Ortholog coalescent tree
4. Ortholog concatenated tree
5. Results of WGD mapping on ASTRAL-Pro tree

**Supplementary Material 1** List of accessions included in this study, including the form of the sample, sample ID used in this study, type of material used for DNA extraction ('source'; H=herbarium, S=silica-dried, SRA=GenBank's SRA), collector and collector number of the sample and voucher for the herbarium specimen, and SRA accession number for the data on NCBI's GenBank. Name changes of samples following this study are in bold under 'Revised name'.

| Sample name | Form | Sample ID | Source | Collector | Collector no. | Herbarium voucher | SRA accession | Revised name |
|---|---|---|---|---|---|---|---|---|
| *Capparis anomala* | - | G09670 | H | Williams | E3P5 | AQ522470 | SRR35419600 | *Capparis anomala* |
| *Capparis arborea* 1 | Northern | G09672 | H | Ford | 2304 | QRS119603 | SRR35419599 | *Capparis arborea* |
| *Capparis arborea* 2 | Northern | G09625 | S | Cooper | WWC2744 | CNS152655 | SRR35419588 | *Capparis arborea* |
| *Capparis arborea* 3 | Southern | G09673 | H | Forster | PIF12587 | QRS106363 | SRR35419577 | *Capparis arborea* |
| *Capparis batianoffii* | - | G09674 | H | Batianoff & Figg | 940401Z | AQ626700 | SRR35419566 | *Capparis batianoffii* |
| *Capparis canescens* | - | G09626 | S | Cooper | WWC2625 | CNS151016 | SRR35419561 | *Capparis canescens* |
| *Capparis humistrata* | - | G09678 | H | Lokkers | s.n. | AQ799027 | SRR35419560 | *Capparis humistrata* |
| *Capparis jacobsii* | - | G09679 | H | Mitchell | 8671 | AQ840980 | SRR35419559 | *Capparis jacobsii* |
| *Capparis lanceolaris* | - | G09680 | H | Ford | 5775 | CNS134441 | SRR35419558 | *Capparis lanceolaris* |
| *Capparis lasiantha* | - | G09681 | H | Gandini | 1121 | CNS133338 | SRR35419557 | *Capparis lasiantha* |
| *Capparis loranthifolia* var. *bancroftii* | - | G09682 | H | Bush & Halford | JJH557 | AQ101020 | SRR35419598 | ***Capparis bancroftii*** |
| *Capparis loranthifolia* var. *loranthifolia* | - | G09683 | H | McLennan | BM07218-3 | AQ1009182 | SRR35419597 | ***Capparis loranthifolia*** |
| *Capparis lucida 1* | *lucida s.s.* | G09684 | H | Cooper | WWC2638 | CNS152670 | SRR35419596 | *Capparis lucida* |
| *Capparis lucida 2* | *aff. arborea* | G09671 | H | Cooper | WWC2415 | CNS145648.1 | SRR35419595 | *Capparis lucida* |
| *Capparis mitchellii* | - | G09628 | S | Jensen, R. | 2344A | AQ0852574 | SRR35419594 | *Capparis mitchellii* |
| *Capparis nobilis* | - | G09685 | H | Sattler & Ziesing | 17 | CBG9708368 | SRR35419593 | *Capparis nobilis* |
| *Capparis ornans* | - | G09692 | H | Forster & Tucker | PIF29872 | AQ577539 | SRR35419592 | *Capparis ornans* |
| *Capparis quiniflora* | - | G09631 | S | Cooper | WWC2600 | CNS151149 | SRR35419591 | *Capparis quiniflora* |
| *Capparis sarmentosa* | - | G09693 | H | Forster | 13884 | QRS104580 | SRR35419590 | *Capparis sarmentosa* |
| *Capparis sepiaria* | - | G08822 | S | Cooper | WWC2615 | CNS151013 | SRR35419589 | *Capparis sepiaria* |
| *Capparis shanesiana* | - | G09694 | H | Cumming | 24773 | CNS146199 | SRR35419587 | *Capparis shanesiana* |
| *Capparis* sp. Bamaga (V.Scarth-Johnson 1048A) | | G08820 | S | Cooper | WWC2585 | CNS151465 | SRR35419586 | ***Capparis xylocarpa*** |
| *Capparis* sp. Coen (L.S.Smith 11862) | - | G09632 | S | Cooper | WWC2362 | CNS144548 | SRR35419585 | ***Capparis megacarpa*** |
| *Capparis* sp. hairy-leaved form | - | LEMJ344 | H | Forster | PIF33222 | AQ0743409 | SRR35419584 | ***Capparis loxophleba*** |
| *Capparis* sp. Wet Tropics vine | - | G08821 | S | Cooper | WWC2621 | CNS152672 | SRR35419583 | ***Capparis splendidissima*** |

| | | | | | | | | |
|---|---|---|---|---|---|---|---|---|
| *Capparis spinosa* subsp. *cordifolia* | Vanuatu | G09667 | H | | | AQ595745 | SRR35419582 | *Capparis spinosa* subsp. *cordifolia* |
| *Capparis spinosa* subsp. *nummularia 1* | WA | G09691 | H | McKenzie | s.n. | PERTH01264001 | SRR35419581 | ***Capparis spinosa* subsp. *nummularia*** |
| *Capparis spinosa* subsp. *nummularia 2* | WA | G09689 | H | Eichler | 22575 | CANB417718 | SRR35419580 | ***Capparis spinosa* subsp. *nummularia*** |
| *Capparis spinosa* subsp. *nummularia 3* | WA | G09688 | H | Smithson | s.n. | CBG7906350 | SRR35419579 | ***Capparis spinosa* subsp. *nummularia*** |
| *Capparis spinosa* subsp. *nummularia 4* | WA | G09690 | H | Lascelles | s.n. | CANB278033 | SRR35419578 | ***Capparis spinosa* subsp. *nummularia*** |
| *Capparis spinosa* subsp. *nummularia 5* | E. QLD | G09677 | H | Specht | LI 353 | AQ028353 | SRR35419576 | ***Capparis spinosa* subsp. *insularis*** |
| *Capparis spinosa* subsp. *nummularia 6* | E. QLD | G09676 | H | Stoddart | 4642 | BRI181370 | SRR35419575 | ***Capparis spinosa* subsp. *insularis*** |
| *Capparis spinosa* subsp. *nummularia 7* | E. QLD | G09627 | S | Cooper | WWC2577 | CNS152674 | SRR35419574 | ***Capparis spinosa* subsp. *insularis*** |
| *Capparis spinosa* subsp. *nummularia 8* | E. QLD | G09675 | H | Cooper | WWC2733 | CNS155748 | SRR35419573 | ***Capparis spinosa* subsp. *formicosa*** |
| *Capparis spinosa* subsp. *nummularia 9* | NT | LEMJ341 | H | Chippendale | 778 | AQ028365 | SRR35419572 | ***Capparis spinosa* subsp. *formicosa*** |
| *Capparis spinosa* subsp. *nummularia 10* | NT | LEMJ343 | H | Perry | 3508 | AQ028371 | SRR35419571 | ***Capparis spinosa* subsp. *formicosa*** |
| *Capparis spinosa* subsp. *nummularia 11* | QLD | G09669 | H | S. Barr | 72 | AQ664543 | SRR35419570 | ***Capparis spinosa* subsp. *formicosa*** |
| *Capparis spinosa* subsp. *nummularia 12* | QLD | G09687 | H | Sankowsky | 3261 | AQ788146 | SRR35419569 | ***Capparis spinosa* subsp. *formicosa*** |
| *Capparis spinosa* subsp. *nummularia 13* | QLD | G09686 | H | Thomas | 3148 | AQ0840672 | SRR35419568 | ***Capparis spinosa* subsp. *formicosa*** |
| *Capparis spinosa* subsp. *nummularia 14* | QLD | G09629 | S | Cooper | WWC2643 | CNS152659 | SRR35419567 | ***Capparis spinosa* subsp. *formicosa*** |
| *Capparis spinosa* subsp. *nummularia 15* | QLD | G09630 | S | Cooper | WWC2748 | CNS155323 | SRR35419565 | ***Capparis spinosa* subsp. *formicosa*** |
| *Capparis thozetiana* | - | LEMJ340 | H | Batianoff | 91101 | QRS107341 | SRR35419564 | *Capparis thozetiana* |
| *Capparis umbonata* | - | LEMJ337 | H | Lothian | 451 | M | SRR35419563 | *Capparis umbonata* |
| *Capparis velutina* | - | G09696 | H | Forster | PIF18279 | QRS116401 | SRR35419562 | *Capparis velutina* |
| *Buchholzia coriacea* | - | - | SRA | Merella *et al.* | 1656 | M05076734 | ERR7621932 | *Buchholzia coriacea* |
| *Cadaba capparoides* | - | - | SRA | Sankowsky | 2256 | PERTH6863361 | ERR7599206 | *Cadaba capparoides* |
| *Maerua glauca* | | - | SRA | Rodin | 4487 | US1991472 | ERR7622173 | *Maerua glauca* |
| *Thilachium africanum* | | - | SRA | Hall | 251 | - | ERR7621968 | *Thilachium africanum* |
| *Boscia albitrunca* | | - | SRA | Kitchin | 63 | NY03946983 | ERR7622164 | *Boscia albitrunca* |

**Supplementary Material 2** Table of loci extraction and HybPhaser statistics for our dataset. CAPTUS statistics are for loci that were used for species tree inference; HybPiper statistics were for loci were used for the calculations of allele divergence, heterozygosity and proportion of Single Nucleotide Polymorphisms (SNPs).

| Sample | CAPTUS | | | HybPiper | | HybPhaser | | | | |
|---|---|---|---|---|---|---|---|---|---|---|
| | No. loci recovered | Recovery (%) | No. copies per locus | No. loci recovered | Recovery (%) | Allele divergence (%) | Heterozygosity | Loci with >0.5% SNPs (%) | Loci with >1% SNPs (%) | Loci with >2% SNPs (%) |
| *Boscia albitrunca* | 347 | 83.44 | 1.28 | 315 | 83.3 | 0.95 | 82.85 | 48.55 | 26.16 | 11.05 |
| *Buchholzia coriacea* | 352 | 89.60 | 1.11 | 322 | 86.9 | 0.32 | 43.57 | 10.53 | 6.14 | 4.09 |
| *Cadaba capparoides* | 351 | 86.66 | 1.22 | 317 | 84.3 | 0.42 | 61.88 | 19.65 | 7.92 | 5.57 |
| *Capparis anomala* | 346 | 81.29 | 1.88 | 309 | 81.5 | 2.23 | 89.43 | 66.77 | 47.13 | 29.91 |
| *Capparis arborea* 1 | 347 | 75.17 | 2.36 | 310 | 81.4 | 2.36 | 93.22 | 76.99 | 57.23 | 35.1 |
| *Capparis arborea* 2 | 349 | 76.08 | 3.47 | 316 | 85.4 | 2.53 | 94.43 | 81.23 | 59.82 | 34.31 |
| *Capparis arborea* 3 | 342 | 74.37 | 1.88 | 320 | 79.4 | 2.22 | 87.8 | 72.32 | 52.98 | 34.52 |
| *Capparis batianoffii* | 349 | 87.79 | 1.36 | 302 | 84.1 | 1.10 | 77.55 | 40.23 | 24.2 | 13.7 |
| *Capparis canescens* | 347 | 83.95 | 1.65 | 326 | 82.9 | 2.03 | 79.22 | 54.82 | 41.27 | 30.12 |
| *Capparis humistrata* | 348 | 80.73 | 2.04 | 301 | 84.9 | 2.34 | 88.82 | 69.71 | 46.76 | 30.88 |
| *Capparis jacobsii* | 351 | 76.08 | 2.93 | 320 | 84.2 | 2.49 | 91.47 | 77.35 | 60.88 | 37.06 |
| *Capparis lanceolaris* | 345 | 82.04 | 1.36 | 322 | 80.4 | 1.72 | 58.21 | 36.42 | 33.13 | 28.06 |
| *Capparis lasiantha* | 346 | 86.70 | 1.24 | 335 | 82.5 | 1.07 | 70.18 | 32.75 | 21.35 | 12.57 |
| *Capparis loranthifolia* var. *bancroftii* | 350 | 84.67 | 1.64 | 305 | 84.6 | 1.99 | 73.24 | 48.53 | 37.35 | 28.53 |
| *Capparis loranthifolia* var. *loranthifolia* | 349 | 78.29 | 2.89 | 333 | 86.3 | 2.45 | 90.67 | 77.26 | 55.39 | 34.69 |
| *Capparis lucida* | 349 | 84.73 | 1.72 | 318 | 85.4 | 1.95 | 69.53 | 46.75 | 38.17 | 26.63 |
| *Capparis lucida aff. arborea* | 348 | 82.91 | 1.61 | 326 | 82.8 | 2.08 | 75.6 | 56.93 | 41.57 | 31.02 |
| *Capparis mitchellii* | 352 | 83.48 | 1.91 | 335 | 85.5 | 2.12 | 80.65 | 57.18 | 39.88 | 28.74 |
| *Capparis nobilis* | 348 | 84.52 | 1.45 | 331 | 83.2 | 1.75 | 57.65 | 40.29 | 33.53 | 27.94 |
| *Capparis ornans* | 346 | 81.83 | 1.46 | 294 | 82 | 1.81 | 74.11 | 47.92 | 38.39 | 29.46 |
| *Capparis quiniflora* | 348 | 88.99 | 1.25 | 326 | 83.2 | 1.01 | 57.27 | 28.49 | 19.29 | 14.24 |
| *Capparis sarmentosa* | 341 | 80.59 | 1.27 | 310 | 76.7 | 0.99 | 56.55 | 32.74 | 21.13 | 13.39 |
| *Capparis sepiaria* | 348 | 86.68 | 1.46 | 309 | 84.7 | 1.51 | 60.88 | 36.18 | 31.76 | 24.71 |
| *Capparis shanesiana* | 345 | 70.37 | 4.12 | 337 | 81.7 | 2.44 | 94.96 | 78.64 | 59.64 | 35.61 |
| *Capparis* sp. Bamaga (V.Scarth-Johnson 1048A) | 348 | 89.68 | 1.26 | 89 | 84.9 | 1.00 | 49.85 | 25.66 | 19.53 | 14.29 |
| *Capparis* sp. Coen (L.S.Smith 11862) | 349 | 86.16 | 1.53 | 172 | 84.3 | 1.96 | 61.54 | 42.31 | 36.09 | 30.18 |
| *Capparis* sp. hairy-leaved form | 351 | 83.76 | 1.40 | 315 | 27.1 | 0.47 | 30.1 | 23.3 | 13.59 | 8.74 |
| *Capparis* sp. Wet Tropics vine | 349 | 71.88 | 5.19 | 336 | 83.3 | 2.67 | 94.01 | 82.04 | 64.67 | 37.43 |
| *Capparis spinosa* subsp. *cordifolia* | 345 | 73.57 | 1.63 | 299 | 75.6 | 1.86 | 92.47 | 83.13 | 68.98 | 25.6 |
| *Capparis spinosa* subsp. *nummularia 1* | 328 | 70.18 | 1.10 | 295 | 65 | 0.59 | 49.84 | 24.6 | 14.38 | 7.67 |
| *Capparis spinosa* subsp. *nummularia 2* | 351 | 88.64 | 1.53 | 302 | 88.1 | 0.92 | 81.32 | 39.37 | 19.83 | 10.06 |
| *Capparis spinosa* subsp. *nummularia 3* | 346 | 86.24 | 1.33 | 306 | 81.9 | 0.94 | 83.38 | 47.18 | 19.29 | 9.79 |

| | | | | | | | | | | |
|---|---|---|---|---|---|---|---|---|---|---|
| *Capparis spinosa* subsp. *nummularia 4* | 342 | 84.47 | 1.18 | 313 | 78.4 | 0.80 | 74.47 | 32.22 | 17.63 | 10.03 |
| *Capparis spinosa* subsp. *nummularia 5* | 343 | 81.11 | 1.11 | 306 | 74.6 | 0.52 | 41.85 | 13.23 | 9.54 | 7.38 |
| *Capparis spinosa* subsp. *nummularia 6* | 335 | 72.54 | 1.48 | 318 | 68.4 | 2.24 | 78.48 | 68.99 | 61.08 | 43.99 |
| *Capparis spinosa* subsp. *nummularia 7* | 349 | 91.03 | 1.16 | 316 | 85.3 | 0.52 | 35.59 | 12.35 | 9.12 | 7.65 |
| *Capparis spinosa* subsp. *nummularia 8* | 350 | 91.25 | 1.17 | 51 | 82.2 | 0.49 | 27.11 | 10.24 | 9.04 | 7.53 |
| *Capparis spinosa* subsp. *nummularia 9* | 347 | 80.93 | 1.12 | 319 | 14.1 | 0.05 | 6.15 | 3.08 | 3.08 | 1.54 |
| *Capparis spinosa* subsp. *nummularia 10* | 350 | 90.30 | 1.14 | 65 | 43.5 | 0.11 | 10.98 | 6.36 | 3.47 | 2.31 |
| *Capparis spinosa* subsp. *nummularia 11* | 347 | 83.91 | 1.13 | 313 | 79 | 0.54 | 49.4 | 18.07 | 11.45 | 8.73 |
| *Capparis spinosa* subsp. *nummularia 12* | 342 | 84.63 | 1.17 | 310 | 79.2 | 0.52 | 52.4 | 17.66 | 10.78 | 7.49 |
| *Capparis spinosa* subsp. *nummularia 13* | 350 | 87.10 | 1.15 | 302 | 81.9 | 0.59 | 48.66 | 17.51 | 10.98 | 7.72 |
| *Capparis spinosa* subsp. *nummularia 14* | 346 | 91.00 | 1.21 | 338 | 83.2 | 0.55 | 39.21 | 15.2 | 10.33 | 7.9 |
| *Capparis spinosa* subsp. *nummularia 15* | 348 | 90.71 | 1.21 | 307 | 84.5 | 0.57 | 51.91 | 16.42 | 10.26 | 8.21 |
| *Capparis thozetiana* | 199 | 51.54 | 1.11 | 101 | 2 | 0.36 | 15.38 | 15.38 | 15.38 | 7.69 |
| *Capparis umbonata* | 347 | 79.10 | 1.36 | 13 | 21.2 | 0.28 | 21.98 | 13.19 | 8.79 | 4.4 |
| *Capparis velutina* | 275 | 56.59 | 1.23 | 62 | 46.7 | 1.29 | 56.44 | 38.64 | 31.06 | 22.73 |
| *Maerua glauca* | 334 | 69.36 | 1.15 | 261 | 70.8 | 0.87 | 64.37 | 40.42 | 25.45 | 11.68 |
| *Thilachium africanum* | 351 | 81.33 | 2.21 | 324 | 89.2 | 1.66 | 97.09 | 83.72 | 57.27 | 17.15 |

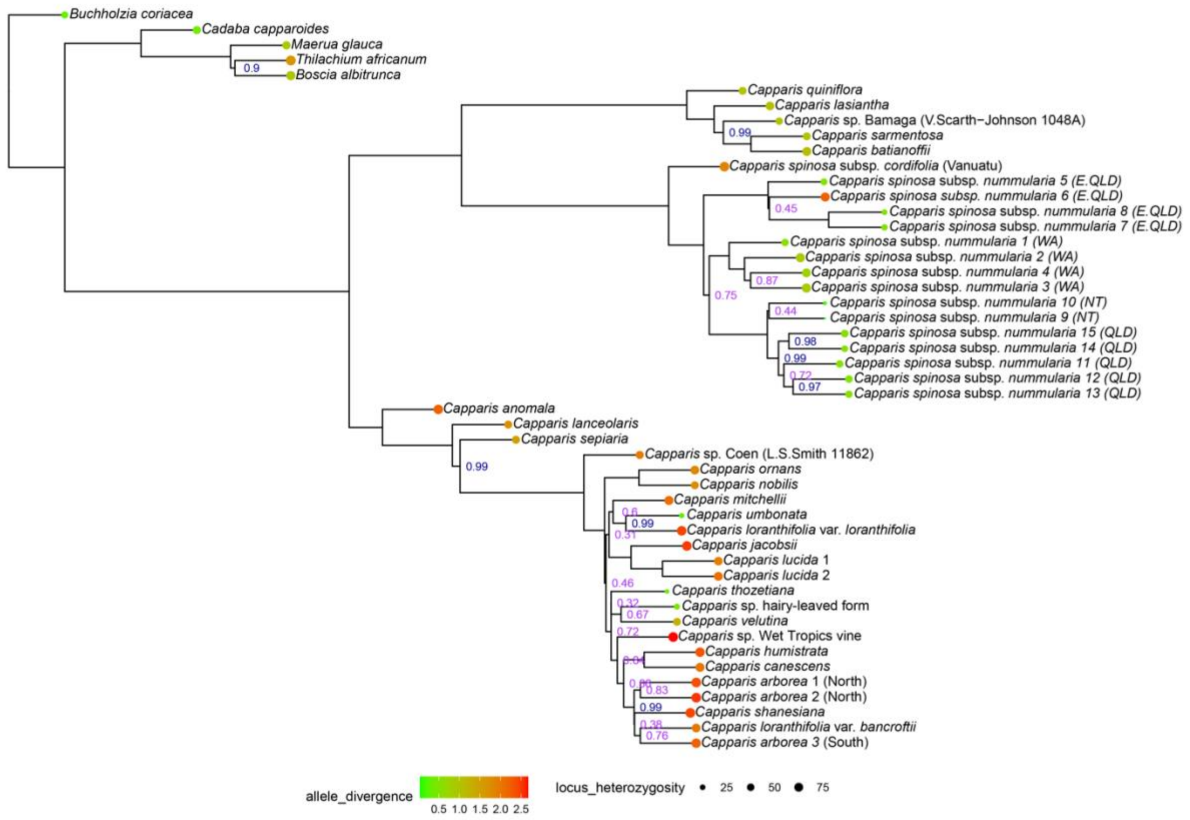


**Supplementary Material 3** Coalescent ortholog tree with locus heterozygosity and allele divergence as estimated by HybPhaser on tips. Posterior probability (PP) indicated on nodes; nodes without a PP value indicated received full support (PP=1).

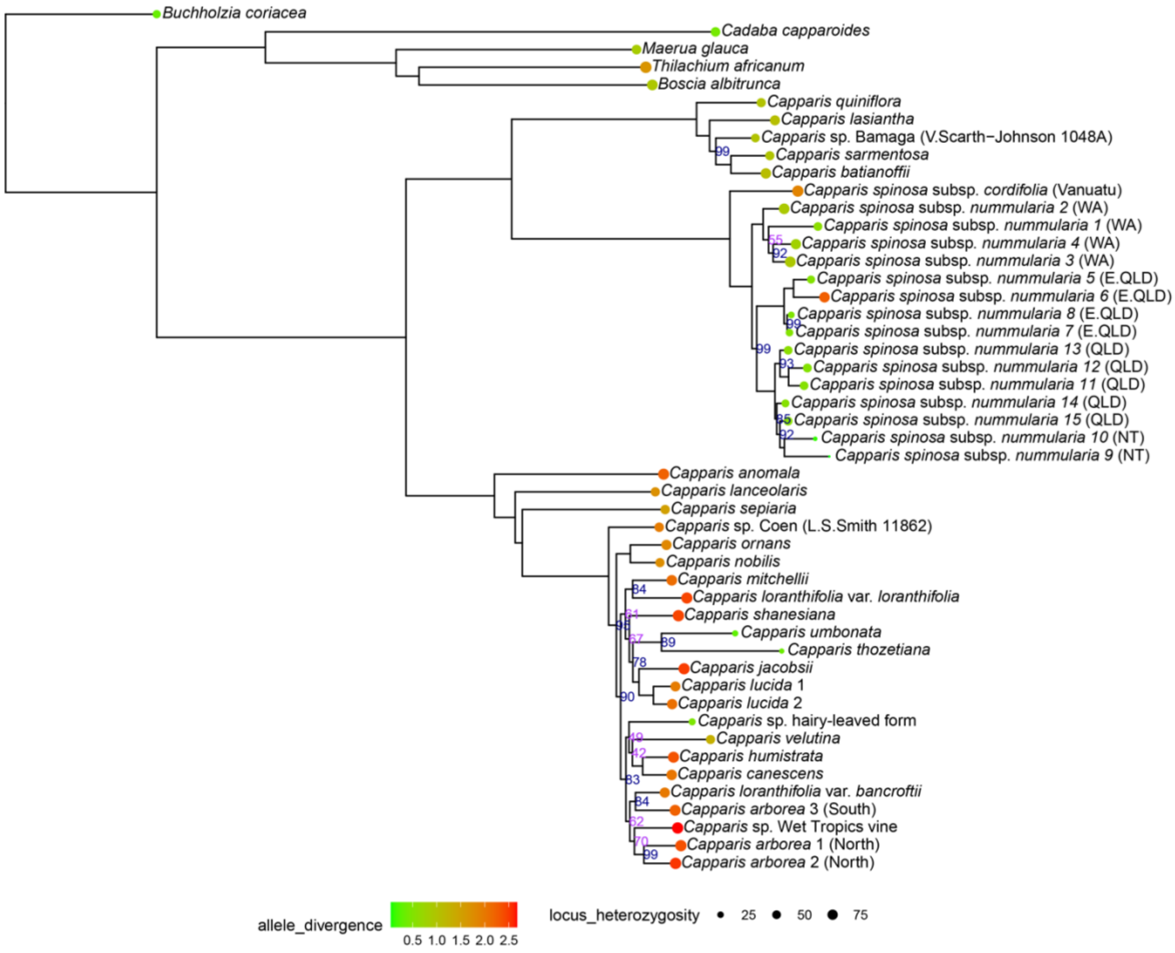


**Supplementary Material 4** Concatenated ortholog tree with locus heterozygosity and allele divergence as estimated by HybPhaser on tips. Bootstrap support (BS) indicated on nodes; nodes without a BS value indicated received full support (BS=100).

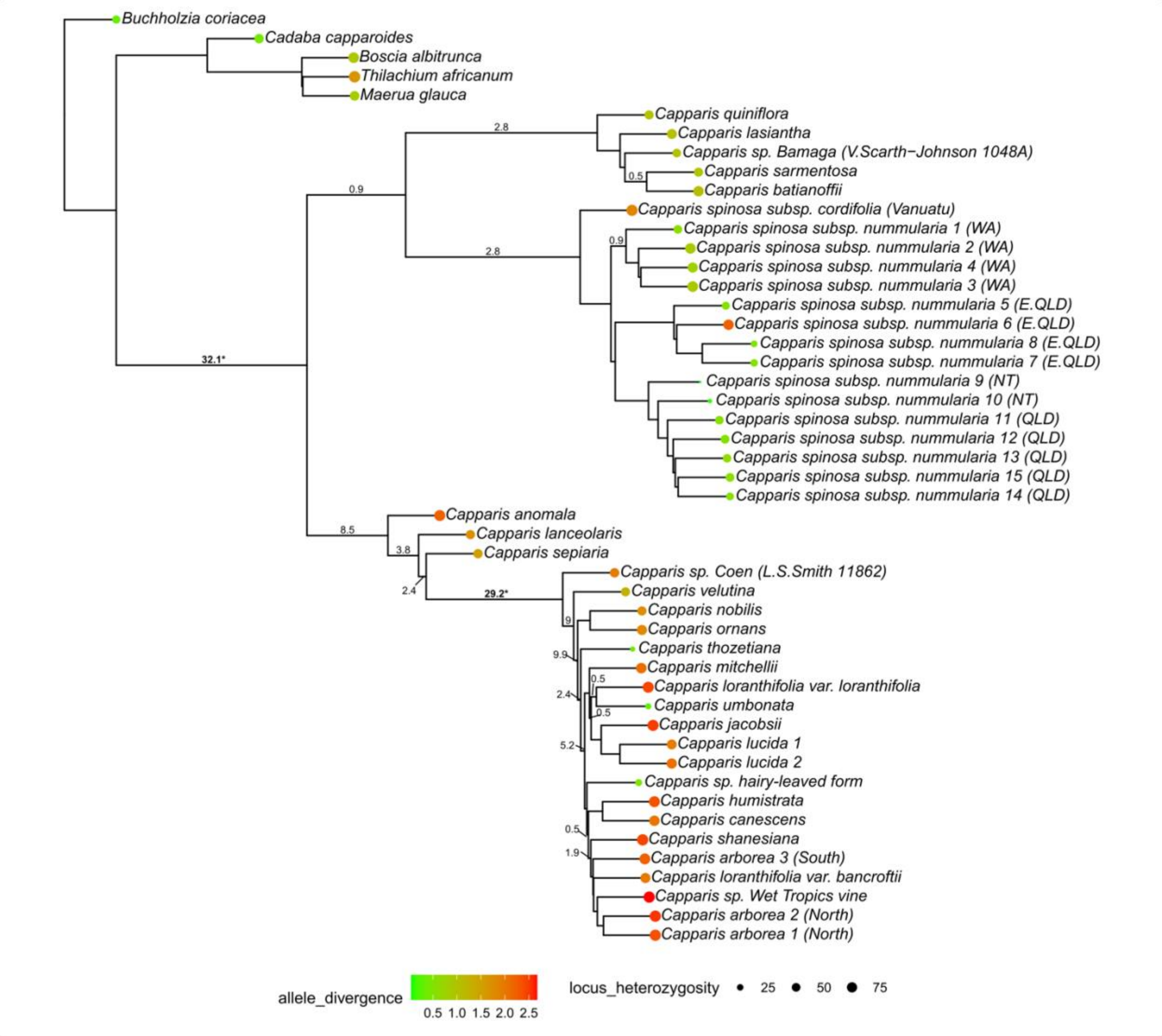


**Supplementary Material 5** Results of WGD mapping analysis on paralog-informed species tree estimated with ASTRAL-Pro. Bolded values with asterisks indicate outlying values, indicating a high likelihood of a WGD event at that node.
